# scPharm: identifying pharmacological subpopulations of single cells for precision medicine in cancers

**DOI:** 10.1101/2023.12.11.571182

**Authors:** Peng Tian, Jie Zheng, Yue Xu, Tao Wu, Shuting Chen, Yinuo Zhang, Bingyue Zhang, Keying Qiao, Yuxiao Fan, Chiara Ambrogio, Haiyun Wang

## Abstract

Intratumour heterogeneity is a major challenge that limits the effectiveness of anticancer therapies, thus compromising treatment outcomes. Single-cell RNA sequencing (scRNA-seq) technology offers a means to capture gene expression profiles at a single-cell resolution, while drug perturbation experiments yield valuable pharmacological data at the bulk cell level. Here, we introduce “scPharm”, a computational framework to integrate large-scale pharmacogenomics profiles with scRNA-seq data, for identifying pharmacological subpopulations within a tumour and prioritizing tailored drugs. scPharm assesses the distribution of the identity genes of single cell (Cell-ID) within drug response-determined gene list, which is accomplished using the normalized enrichment score (NES) obtained from Gene Set Enrichment Analysis (GSEA) as the statistic. One key strength of scPharm is rooted in the robust positive correlation between NES statistics and drug response values at single-cell resolution. scPharm successfully identifies sensitive subpopulations in ER-positive and HER2-positive human breast cancer tissues, discovers dynamic changes in resistant subpopulation of human PC9 cells treated with Erlotinib, and expands its prediction capabilities to a mouse model. By a thoroughly comparative evaluation with other single-cell prediction tools, scPharm presents the superior predictive performance and computational efficiency. Furthermore, scPharm offers a unique feature by predicting combination strategies, gauging compensation effects or booster effects between two drugs through the Set covering method, as well as evaluating drug toxicity on healthy cells within the tumour microenvironment. Together, scPharm provides a novel approach to uncover therapeutic heterogeneity within tumours at single-cell resolution and facilitates precision medicine in cancers.

## Introduction

The intrinsic heterogeneity of individual tumours poses a significant challenge in cancer therapeutics, often leading to the failure of anticancer therapies and the death of patients. Recent advances in single-cell sequencing technology have opened avenues for investigating molecular heterogeneity at the single-cell level, offering promise for precision cancer therapy. In recent years, technologies like Perturb-seq [1], ECCITE-seq [2] and sci-Plex [3] integrate single-cell sequencing with compound or CRISPR-mediated gene perturbation experiments, allowing the assessment of single-cell perturbation response at the transcriptome or surface protein level. Despite their potential, these biotechnological platforms are still in their infancy.

Remarkably, the application of perturbation-based screening in vivo at bulk cells of cancer cell lines over the past decade has proven valuable in identifying cancer cell targets. Moreover, such screening has produced large-scale genetic characterizations of cancer cell lines as well as their phenotype in perturbation experiments. For example, the pharmacogenomic projects Cancer Cell Line Encyclopedia (CCLE) [4] and Genomics of Drug Sensitivity in Cancer (GDSC) [5] performed sequencing of over 1,000 tumour cell lines and sensitivity analysis of hundreds of drugs. Following this, the Cancer Therapeutics Response Portal (CTRP) [6] has tested the drug response of nearly 500 drugs against almost 900 cancer cell lines. In addition, CRISPR screening platform projects with the concept of joint lethality, including Project Score [7], Project Achilles [8], and Project DRIVE [9], identified genetic perturbation required for cell fitness in molecular contexts. Given these advancements, there is potential to develop a computational pipeline that integrates single-cell transcriptomic profiles with pharmacogenomics profiles to predict the pharmacological phenotype of single cells, addressing the limitations of current biotechnology platforms.

While some sporadic work [10, 11] has emerged to predict drug response of single cells by co-embedding expression profiles of single cells and bulk cancer cells using machine learning-based methods, challenges persist. These challenges extend beyond the prediction task itself to interpreting results and gaining insights into precision medicine. Machine learning-based methods face major limitations, including the requirement for large datasets with sufficient heterogeneity for model training. Publicly available pharmacogenomic datasets are still limited in size compared to the complexity of the model. Additionally, these methods often rely on binarized labels of drug response, such as sensitive vs. resistant, introducing a dependence on the binarization method that may lack biological interpretation. Moreover, most existing methods construct models solely to predict whether single cells are sensitive or resistant to drugs, overlooking the broader clinical context of drug efficacy and side effects.

Here we propose “scPharm”, a computational framework appliable to single-cell RNA sequencing (scRNA-seq) data of tumour tissue in the real scenario, aiming to address these challenges. These scenarios involve a mixture of various cell types within the tumour microenvironment, rather than focusing solely on pure cancer cells. We hypothesize that if a single cell is sensitive to a drug, the highly expressed genes in this cell, termed Identity genes (Cell-ID), would significantly overlap with genes associated with increased drug sensitivity. Conversely, if a single cell is resistant, the highly expressed genes would overlap with those correlated with increased drug resistance. Our approach employs Gene Set Enrichment Analysis (GSEA) [12] as a paradigm, dynamically calculating Cell-ID as the gene set based on scRNA-seq data. Unlike regular GSEA, which uses a stable set of a priori defined genes, scPharm utilizes the dynamically calculated identity genes of each single cell. Accordingly, a ranked list of genes is drug response-determined which is generated by the correlation of gene expression with drug response based on bulk data. The gene list per drug within the same cancer type is stable, applicable to all single cells.

In contrast to machine learning models that transfer drug response phenotypes from bulk data to single cells based on expression profile similarity, scPharm utilizes the normalized enrichment score (NES) derived from GSEA to identify statistically significant sensitive and resistant single cells. This statistical methodology overcomes challenges related to small sample size, binarization in machine learning models, and the integration of expression similarity across modalities. scPharm’s performance is comprehensively assessed across various datasets, including scRNA-seq data from ER-positive and HER2-positive breast cancer tissues[13], PC9 cells treated with Erlotinib at different time points[14], and the MMTV-PyMT mouse mammary tumor model[15]. Additionally, a rigorous comparative analysis is conducted to assess the predictive performance of scPharm against other single-cell prediction tools, including scDEAL [10], CaDDReS-Sc [16], Scissor [17] and SeuratCCA [18].

Notably, scPharm goes beyond prediction, providing valuable insights into prioritization of tailed drugs. Its distinctive feature also lies in the assessment of combination strategies, gauging compensation effects or booster effects between two drugs through the Set covering method. Moreover, scPharm evaluates drug side effects by leveraging scRNA-seq data from healthy human tissues, enabling the assessment of drug toxicity on healthy cells within the tumour microenvironment.

## Results

### Overview of scPharm

scPharm is a two-module computational frame designed to integrate large-scale pharmacogenomics profiles with scRNA-seq data (Fig. 1, See Methods). scPharm is applied to scRNA-seq data obtained from real tumour tissue scenario involving a mixture of various cell types within the tumour microenvironment. scPharm first employs CopyKAT [19] to differentiate tumour cells from healthy cells. **Module 1** focuses on identifying phamacological subpopulations of tumour single cells. It takes two inputs: 1) drug response-determined gene list calculated from bulk RNA-seq and corresponding drug response data from GDSC dataset, where genes at the top correlate with drug resistance and those at the bottom with drug sensitivity, and 2) the identity genes of single cell (Cell-ID) extracted using Multiple Correspondence Analysis (MCA) [20]. The module uses the normalized enrichment score (NES) from GSEA to assess the distribution of the Cell-ID within the ranked gene list and to predict drug response of a single cell. By going through all cells in a tumour, we have the distribution of NES statistics to infer pharmacological subpopulations to a drug. **Module 2** evaluates drug suitability using the Dr score which is calculated by the proportions of sensitive and resistant subpopulations and prioritizes the tailored drugs. It further explores potential combinatorial therapies, gauging compensation effects or booster effects between two drugs, and evaluates potential drug toxicity on healthy cells within the tumour microenvironment.

**Figure 1.**
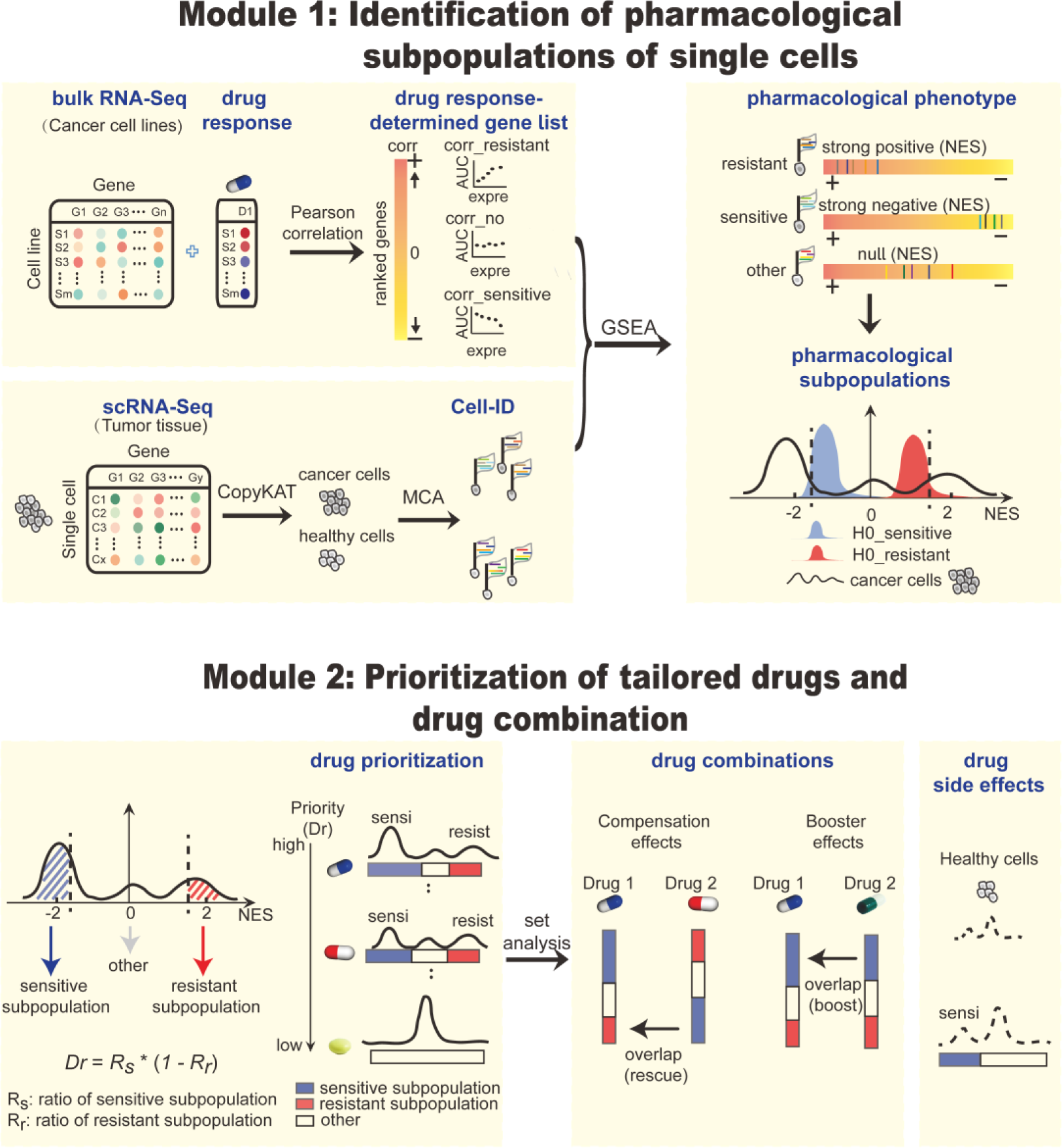
Overview of the computational framework scPharm. scPharm consists of the following two modules. The first module (Module 1) utilizes pharmacogenomic data and scRNA-seq data to identify statistically significant sensitive and resistant single-cell subpopulations. The method Gene Set Enrichment Analysis (GSEA) as a paradigm determines whether a single cell is sensitive to a drug or not. The second module (Module 2) prioritizes tailored drugs by Dr scores and develops drug combination strategies by evaluating whether two drugs have compensation effects or booster effects.

To model a statistical test in Module 1, we utilized scRNA-seq data from healthy human tissues [21-25] and the pooled drugs to generate null distributions (H0) of NES. We investigated three types of healthy human tissues, including lung, breast and skin. There are very similar distribution characteristics among them, featured with two peaks, one with the median around -1 and the other around +1 (Supple. Fig. 1, Fig. 1). We thus pooled three types of healthy tissues and all drugs to generate a unique null distribution. Gaussian mixture model decomposition (See methods) [26] was then performed to accurately estimate the parameter of two separate H0, respectively for sensitive cells (H0_sensitive) and resistant cells (H0_resistant). Using a unified threshold based on the mean plus or minus one standard deviation, respectively for H0_sensitive and H0_resistant, scPharm categorizes tumour single cells into sensitive subpopulation or resistant subpopulation for a given drug.

### Rationale of scPharm

scPharm allows users to predict drug response using NES statistics. We hypothesize that if a single cell is sensitive to a drug, its Cell-ID would significantly overlap with genes whose high expression correlates with increased drug sensitivity (sensitive markers); conversely, if a single cell is resistant to a drug, its Cell-ID would significantly overlap with those correlated with increased drug resistance (resistant markers). To address this issue, GSEA is employed as a suitable paradigm. Our study created drug response-determined gene list using the cell lines within the same cancer type, for the aims to mitigate the influence of tissue-specific expression on predictions.

To test these hypotheses, we first observed the correlation between NES statistic and drug response at the single-cell level. The scRNA-seq data of human cancer cell lines and drug response (AUC) data which is quantified with Area Under the dose response Curve (AUC) from the cell population of the same cancer cell lines were collected (see Methods) [27]. We employed GSEA to compute NES statistics of single cells, then calculated the Pearson correlation between NES and AUC. Our analysis covered 281 drugs and about 40 lung adenocarcinoma (LUAD) cell lines. Remarkably, we observed a very high correlation (R=0.98) for drugs like Docetaxel (Fig. 2A), where 13 LUAD cell lines with each cell line including 252 single cells were investigated. We used the median of NES statistics of these single cells as the summarized NES for the corresponding cell line. This robust correlation extended across 281 drugs, with the median of correlations around 0.8 (Fig. 2B) and other cancers spanning 12 cancer types (Fig. 2C).

**Figure 2.**
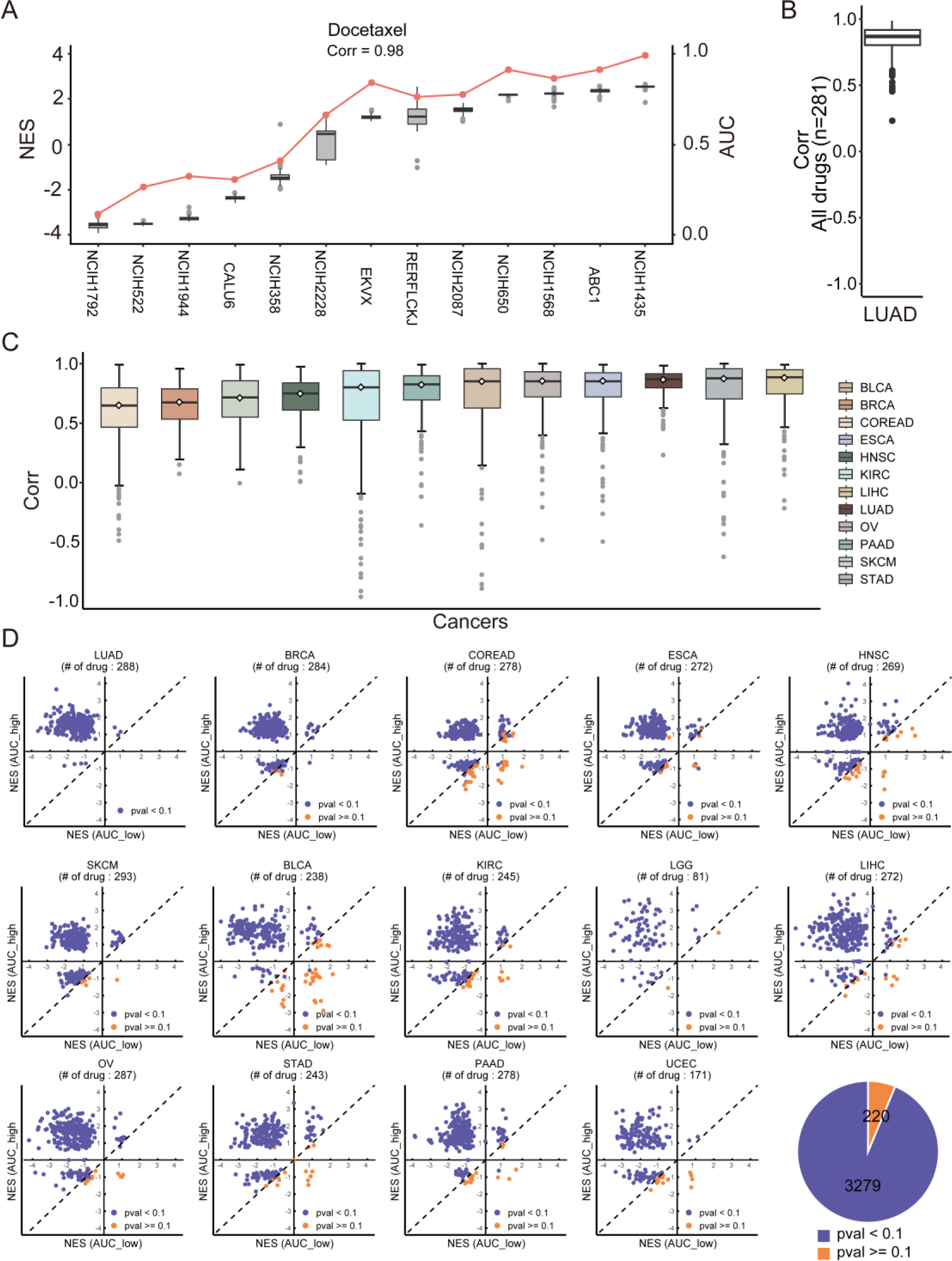
The correlation between NES and drug response. **A.** The red line depicting drug response of 13 lung adenocarcinoma (LUAD) cell lines to Docetaxel, which is measured using the dose-response curve (AUC). The boxplot depicting NES statistics for single cells in the corresponding LUAD cell lines. The Pearson correlation coefficient (Corr) between AUC and NES median of single cells is shown. **B.** The boxplot of Pearson correlation coefficients between AUC and NES median for all drugs in LUAD. **C.** The boxplot of Pearson correlation coefficients across the different cancers. **D.** Scatter plot showing the NES statistics in the AUC_high group versus the AUC_low group. Cell lines are divided into AUC_high and AUC_low groups based on the median of AUC values to each drug. Pie plot depicting the number of drugs with the significant p values regarding the correlations between AUC and NES across the cancers.

Further analysis, dividing single cells into AUC_high vs. AUC_low groups, blue-highlighted statistically higher NES statistics in the AUC_high group for the majority of drugs (Fig. 2D). These analyses using different statistical approaches provide strong evidence that NES is robustly and highly correlated with drug response, demonstrating its predictive potential at the single-cell level.

### Identification of sensitive subpopulation in HER2-positive breast cancer

To assess whether scPharm can accurately identify sensitive subpopulation from single-cell data, we interrogated scRNA-seq data from six HER2-positive human breast cancer tissue samples, named as MH0176, MH0031, MH0069, PM0337, AH0308 and MH0161 [13]. After scRNA-seq data processing, we retained at total of 44,761 cells, with an average of 7,460 cells per sample and 1,356 genes captured per cell. These HER2-positive samples are known to be sensitive to HER2 inhibitions. This evaluation specifically focused on three types of HER2 inhibitions – Afatinib, Saptinib and Lapatinib. We deciphered 10,124 tumour cells and 34,637 healthy cells from all 44,761 cells using CopyKAT, and extracted 200 identity genes per cell using MCA method.

When Afatinib was considered, the NES distribution curves of single tumour cells were featured with one big negative peak and small bumps typically in MH0176, MH0031, MH0069, and PM0337 (Fig. 3A). Interestingly, for all samples except MH0161, the negative peak was significantly left-shifted against H0_sensitive. This indicates the presence of the sensitive subpopulation within the tumour cells. Conversely, this phenomenon was not observed in adjacent normal cells from the same samples (Fig. 3B). Additionally, such distribution curves were also observed when other HER2-targeted drugs, Sapitinib and Lapatinib, were considered (Supple. Fig. 2).

**Figure 3.**
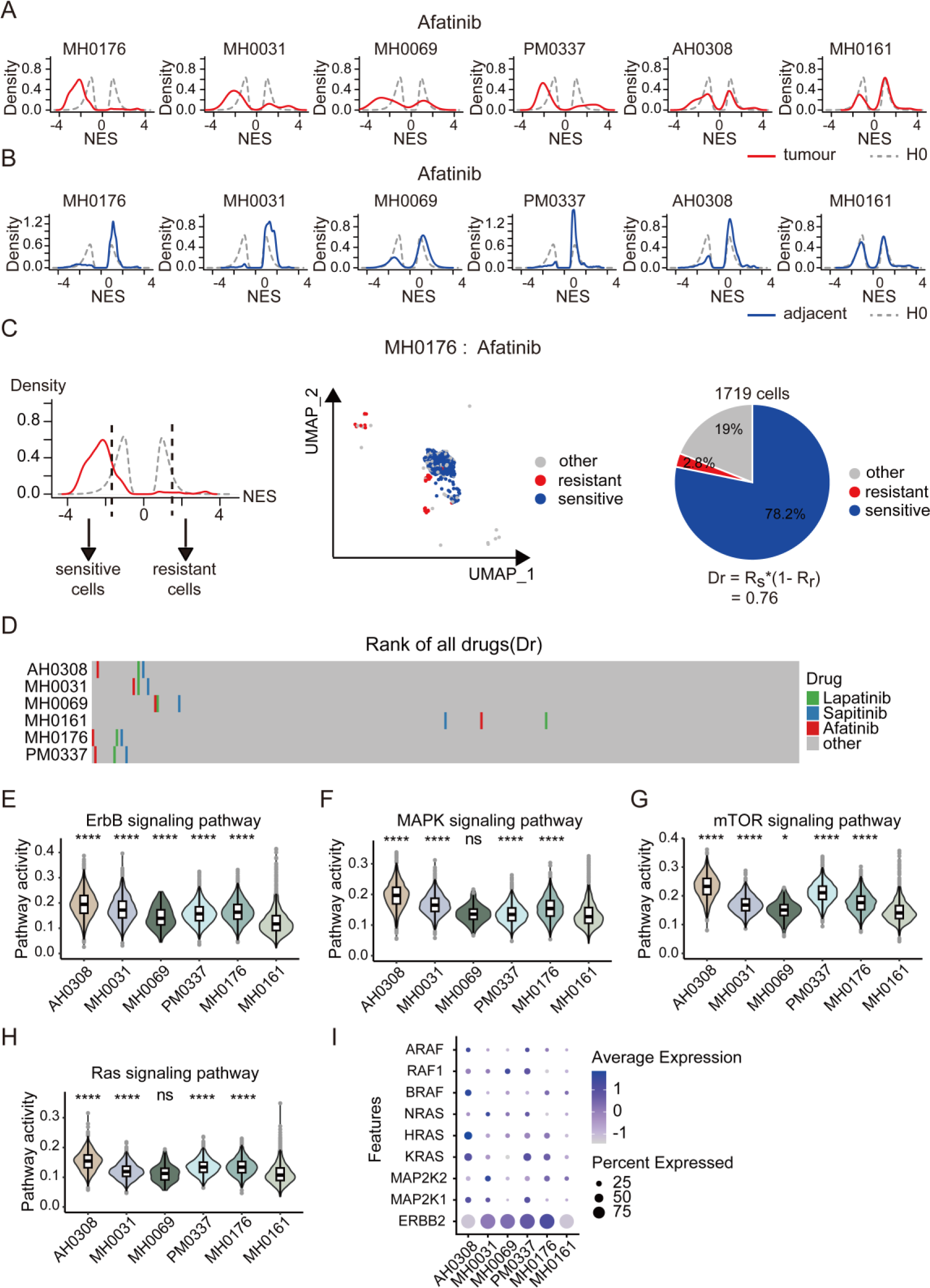
scPharm is applied in HER2+ breast cancer. **A.** Density plot depicting the NES statistics of single cells from six HER2+ breast cancer tissues (red curves) and from healthy human tissues (grey curves) in the context of Afatinib. **B.** Density plot depicting the NES statistics of single cells from tumour-adjacent tissues and from healthy human tissues in the context of Afatinib. **C.** Plot illustrating the sample MH0176 in the context of Afatinib. **D.** Rank of three HER2 inhibitions in HER2+ samples. **E.** ErbB signaling pathway activity comparison for all HER2+ samples. **F.** MAPK signaling pathway activity comparison for all HER2+ samples. **G.** mTOR signaling pathway activity comparison for all HER2+ samples. **H.** Ras signaling pathway activity comparison for all HER2+ samples. **I.** Expression of ERBB2(HER2) and key genes in ErbB signaling pathway. The Wilcoxon rank-sum test was employed between sample MH0161 and other samples, with “ns” indicating a p-value greater than 0.05, “*” indicating a p-value less than 0.05, “**” indicating a p-value less than 0.01, “***” indicating a p-value less than 0.001, and “****” indicating a p-value less than 0.0001.

Based on the null distribution of NES, scPharm utilized the cutoff -1.75 and 1.52 to test whether the NES for a single cell is statistically significant from H0. Then it identified both sensitive and resistant cell subpopulations by going through all cells in a tumour. Given the sample MH0176 in the context of Afatinib for example (Fig. 3C), the sensitive subpopulation includes 1,344 cells identified using cutoff -1.75, and the resistant one includes 49 cells with cutoff 1.52. Based on the cell sizes of two subpopulations, the drug score Dr for Afatinib is 0.76. When retrieving all 295 drugs, we ranked them based their corresponding Dr values, as depicted in Figure 3D. In this figure, three types of HER2 inhibitions were highlighted in green, blue and red colors, with grey color for other drugs. The results showed that among five of six HER2-positive samples, HER2 inhibitions consistently achieved high scores and ranked at the top of all drugs (Fig. 3D). The details of the drugs ranking at the top 30 are shown in Supplemental Figure 3. Notably, drugs targeting EGFR, such as Gefitinib, Erlotinib, and Osimertinib, also ranked at the top, suggesting a potential role for EGFR-targeted drugs in HER2-positive patients[28].

Notably, for sample MH0161, three HER2 inhibitions did not rank at the top as we expected. To unravel the potential mechanism leading to this difference, we investigated the activity of pathways associated with HER2 inhibitors as well as the expression of key genes in these pathways. The pathway activity is measured using the average expression of genes in this pathway. The analyses revealed that the activity of ErbB signaling pathway - a target of HER2 inhibitions - was significantly lower in MH0161 compared to the other five samples (Fig. 3E). Its downstream pathways, such as MAPK signaling, mTOR signaling, and Ras signaling, also exhibited lower activity in MH0161 (Fig. 3F-3H). Moreover, key genes such as ERBB2, KRAS, and MAP2K1 were almost not expressed in MH0161 (Fig. 3I). These findings may explain the discrepancy in the rank of HER2 inhibitions in MH0161.

### Identification of sensitive subpopulation in ER-positive breast cancer

We further evaluated scPharm on scRNA-seq data from 14 ER-positive breast cancer tissue samples [13]. The analyses of scRNA-seq data retained at total of 74,878 cells, with an average of 5,348 cells per sample and 1,041 genes captured per cell. These ER-positive samples are known to be sensitive to ER inhibitions. This evaluation focused on two types of ER inhibitions – Fulvestrant and GDC0810. For Fulvestrant, there are two drug response assays in GDSC dataset, corresponding to the different maximum screening concentration. Here we refer to them by Fulvestrant_low and Fulvestrant_high, respectively according to the maximum concentration 1µm and 10µm. We deciphered 26,251 tumour cells and 48,627 healthy cells from all 74,878 cells and extracted 200 identity genes per cell.

When GDC0810 considered, we robustly observed the left-shifted negative peak in single tumour cells across ER-positive samples with 4 samples were selected to be shown, but not in adjacent healthy cells (Supple. Fig. 4A-B). This indicated the significant existence of sensitive subpopulation to GDC0810. The curves with such characteristic extended to Fulvestrant (Supple. Fig. 5). Based on Dr values of 295 drugs, we ranked them and highlighted ER inhibitions (Supple. Fig. 4C). Except in AH0319 and MH0042, ER inhibitions consistently ranked at the top in the majority of samples (Supple. Fig. 4C). In detail, the top 30 drugs recommended by scPharm in four samples were shown in Supplemental Figure 4D.

Interestingly, when only focusing on Fulvestrant with two maximum screening concentrations, we observed that scPharm ranked Fulvestrant_high ahead of Fulvestrant_low (Supple. Fig. 4C). Since for Fulvestrant_low with 1µm as maximum concentration, drug response values of numerous cell lines are beyond the testing range and cannot be accurately estimated. And the replicated assay using the increased maximum concentration with 10µm can cover the drug response concentration for most of cells to estimate drug response accurately. Accordingly, drug response-determined gene list used for GSEA is more accurate for Fulvestrant_high than Fulvestrant_low. This led to the improved ranking of Fulvestrant_high in ER-positive breast cancer samples (Supple. Fig. 4C, Supple. Fig. 6A).

Further analyses revealed that the sensitive subpopulation to Fulvestrant_high was significantly larger than Fulvestrant_low (Supple. Fig. 6B-G). Based on subpopulation changes between two concentrations, we observed that many cells not efficiently classified in Fulvestrant_low (labelled as other subpopulation) were accurately identified as sensitive one in Fulvestrant_high. This suggests that the accuracy of drug response-determined gene list leads to high efficiency in identification of sensitive and resistant subpopulations, and exhibits scPharm’s diversity in ability to process with pharmacogenomics data with difference in quality.

### Discovery of dynamic changes in resistant subpopulation of PC9 cells treated with Erlotinib

The aforementioned evaluations showcased scPharm’s efficacy in identifying sensitive subpopulations in ER-positive and HER2-positive human breast cancer tissues. To further assess scPharm’s ability to uncover resistant subpopulations and observe their dynamic changes over time, we employed scRNA-seq data from the lung adenocarcinoma cell line PC9. PC9 cells harbor an exon 19 deletion in EGFR gene and were treated with the EGFR inhibitor Erlotinib at Day 1, Day 2, Day 4, and Day 9 [14]. Although PC9 cells are initially sensitive to EGFR inhibition, continuous treatment leads to cells survival and the occurrence of drug resistance at Day 9 and Day 11 (Fig. 4A).

**Figure 4.**
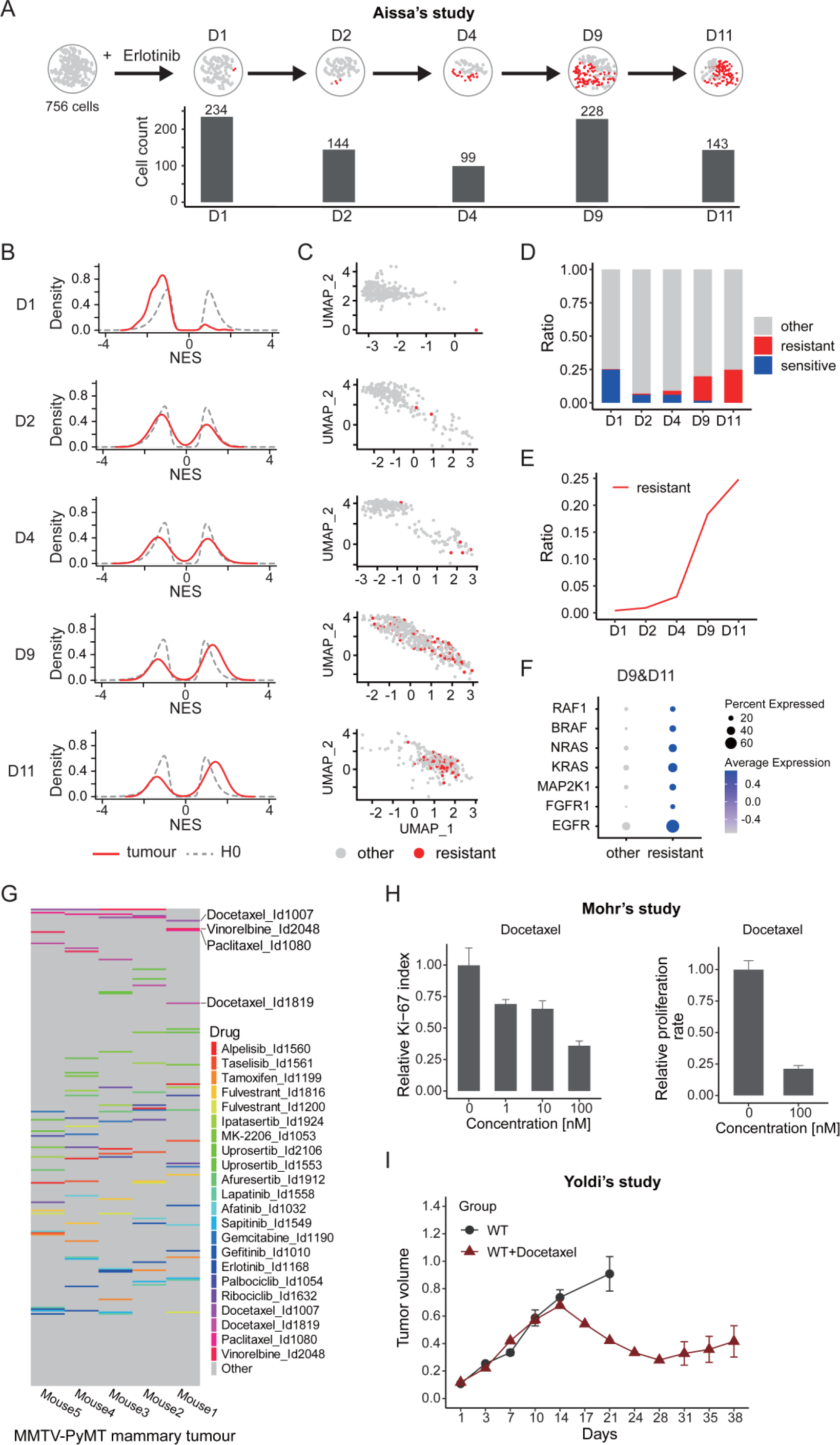
Application of scPharm in PC9 cells and MMTV-PyMT model mice. **A.** Graph showing the number of drug-resistant cells identified by scPharm in PC9 cells treated with Erlotinib at different time points (Days). Temporal cell counts were obtained from the study of Aissa et al. **B.** scRNA-seq data showing the dynamics of the NES peak in PC9 cells. **C.** UMAP visualization highlighting the resistant cells identified by scPharm (depicted in red). **D**. Stacked bar chart depicting the changing proportion of sensitive, resistant and other subpopulations at different time points. **E.** Temporal changes in the proportion change of resistant subpopulation. **F.** Expression of genes in EGFR pathway and downstream pathways. **G.** Heatmap presenting predictive ranking for various clinical therapeutic drugs in breast cancer, derived from Opentarget Platform. The analysis is based on data obtained from the MMTV-PyMT mouse model, encompassing five samples. **H.** Bar graph showing the relative abundance of Ki-67 abundance (left) and relative proliferation rate of breast cancer cells (right) in MMTV-PyMT mice treated with different concentrations of Docetaxel. Data were obtained from the study of Mohr et al. **I.** Diagram showing breast cancer tumour size in MMTV-PyMT mice at different time points (dark red: treated with Docetaxel, grey: control). Data were obtained from the study of Yoldi et al.

This dataset includes a total of 1,311 PC9 cells, with an average of 262 cell at each time point and 1,132 genes captured per cell. The result revealed a rightward shift of the positive peak against H0_resistant at Day 9 and Day 11, indicating the emergence of resistant subpopulation following continuous treatment of Erlotinib (Fig. 4B, C). Notably, at Day 1, the negative peak was left-shifted against H0_sensitive, aligning with the fact that PC9 cells exhibited an initial favorable response to Erlotinib. Tracking the pharmacological subpopulations over time, we observed an increase in resistant subpopulation accompanied by a decline in sensitive subpopulation (Fig. 4D). Particularly noteworthy was the sharp increase in resistant subpopulation after Day 4 (Fig. 4E), precisely coinciding with the experimental findings (Fig. 4A). To understand the mechanisms underlying drug resistance, we further investigated gene expression in EGFR pathway and downstream pathways. We observed the upregulation of EGFR, FGFR1, MAP2K1, KRAS, NRAS, BRAF, and RAF1, indicating the reactivation of signaling downstream of the inhibited EGFR in the resistant PC9 cells (Fig. 4F).

### Expanding scPharm’s prediction capabilities to a mouse model

The mouse is the foremost mammalian model for studying human disease and therapies. While scPharm primarily utilized pharmacogenomic profiles from human cancer cell lines for predicting pharmacological subpopulations, we sought to evaluate its potential applicability to a mouse model. We obtained scRNA-seq data from the well-established MMTV-PyMT mouse mammary tumour model, including samples from five mice [15].

In total, there are 3,504 tumuor single cells, with an average of 700 cells per mouse, and 13,245 orthologous genes in humans per cell included in our evaluation. When ranking drugs based on their Dr scores, we observed that chemotherapeutics such as Docetaxel, Vinorelbine, and Pacilitaxel ranked at the top of the recommended drugs, while targeted drugs did not (Fig. 4G). This aligns with the genetic background of the MMTV-PyMT mouse model, which loses expression of ERα and PR as the tumour progresses and expresses HER2 at a low level [31]. Moreover, a few studies based on in vitro (Fig. 4H) and in vivo (Fig. 4I) experiments have shown that MMTV-PyMT mice are sensitive to Docetaxel treatment, resulting in reduction of tumour cells and tumour shrinkage (Fig. 4I) [29, 30]. Similarly, the report on the sensitivity of Paclitaxel to MMTV-PyMT mouse model also provides favorable evidence for our results [31, 32]. These independent results provide cross-validation of scPharm’s predictive ability when applied to mouse model.

### A comprehensive evaluation of different methods in breast cancer samples

To comprehensively illustrate the predictive performance of scPharm, we conducted a comparative analysis with two widely used single-cell prediction tools, scDEAL [10] and CaDDReS-Sc [16], along with label transfer tools, Scissor[17] and SeuratCCA [18]. Scissor and SeuratCCA are not typically used in this domain.

Evaluation of single-cell prediction models at the individual cell level poses challenges since drug response assays are typically conducted on cell populations rather than individual cells. We then introduced a novel evaluation approach focused on comparing the drug recommendations generated by different tools. ER-positive and HER2-positive breast cancer tissue samples, known to be sensitive to ER or HER inhibitions, were employed for this evaluation.

Initially, sensitive and resistant subpopulations of breast cancer samples were predicted by the different tools. Then we employed Dr value developed by scPharm to rank all drugs in the analysis (See Methods) (Fig. 5A-B). Our focus was on examining the rankings of three HER2 inhibitions Afatinib, Lapatinib, and Sapitinib in HER2-positive breast cancer samples. The findings indicated that, except for sample MH0161, scPharm consistently ranked these drugs at the top and in close proximity (Fig. 5A), significantly outperforming other tools (Fig. 5C). While CaDDReS-Sc exhibited consistent prediction results across different samples, substantial differences were observed between the samples.

**Figure 5.**
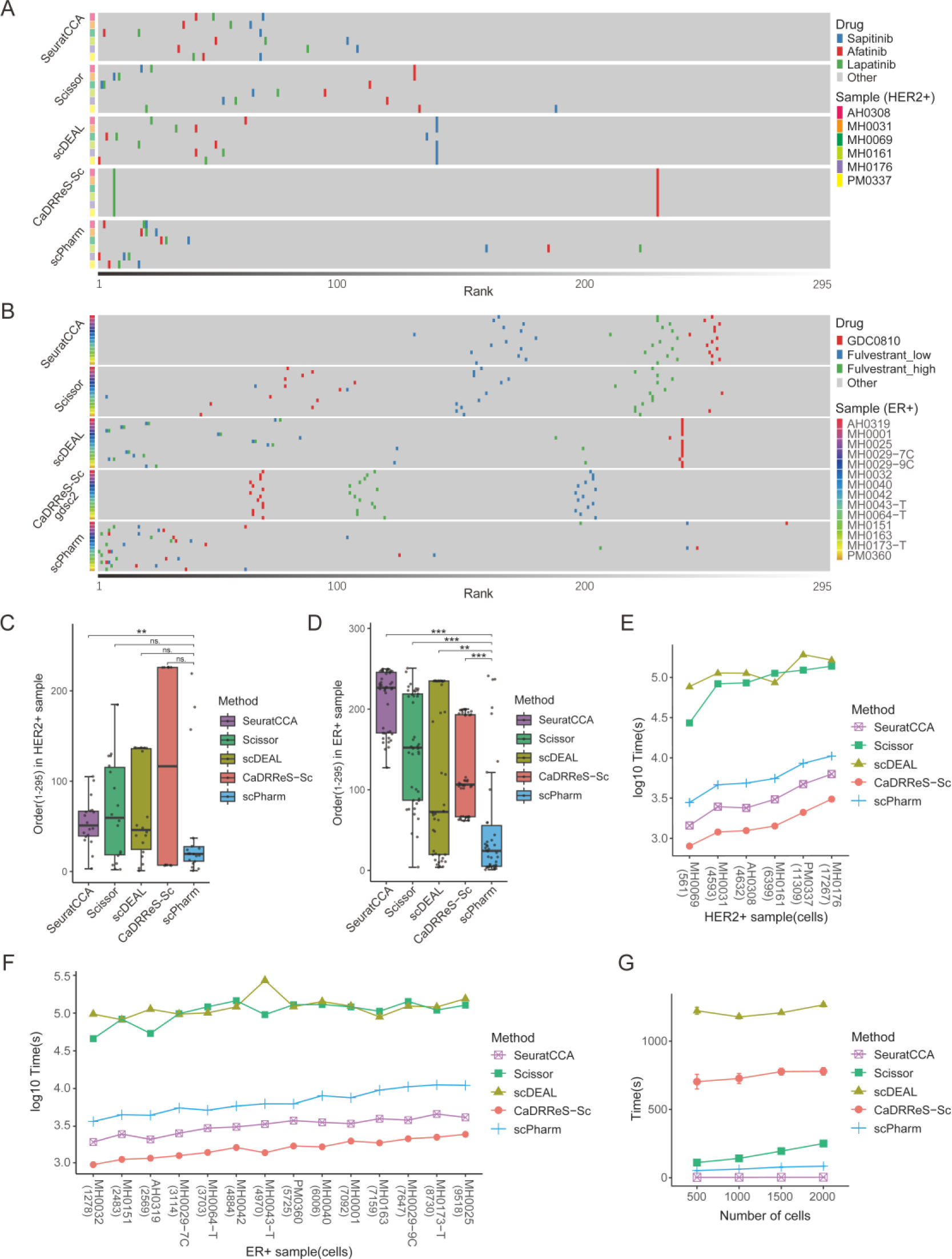
Benchmark of different methods including SeuratCCA, Scissor, scDEAL, CaDRReS-Sc and scPharm. **A.** Predictive ranking of HER2 inhibitions in HER2+ samples across the different methods. **B.** Predictive ranking of ER inhibitions in ER+ samples across the different methods. **C.** Statistical comparison of predictive ranking of HER2 inhibitions in HER2+ samples predicted by the different methods. **D.** Statistical comparison of ranking of ER inhibitions in ER+ samples predicted by the different methods. The Wilcoxon rank-sum test was employed, with “*” indicating a p-value less than 0.05, “**” indicating a p-value less than 0.01, and “***” indicating a p-value less than 0.001. **E.** Comparison of runtime for different methods in HER2+ samples. **F.** Comparison of runtime for different methods in ER+ samples. **G.** Comparison of different methods for calculating runtimes of individual drugs at different single cell numbers sampled from a HER2-positive breast cancer sample (PM0337).

Due to CaDDReS-Sc’s default use of GDSC1 drugs for calculations, the prediction for the drug Sapitinib was missing. To address this, we extended CaDDReS-Sc to utilize GDSC2 data for training and prediction, enabling a comparison of predictions for ER-positive breast cancer samples. In these samples, the rankings of ER inhibitions GDC0810 and Fulvestrant were similarly prominent and concentrated in scPharm (Fig. 5B), significantly surpassing other tools (Fig. 5D).

Subsequently, we assessed the computational time of the tools. Notably, scPharm demonstrated markedly higher efficiency compared to Scissor and scDEAL, albeit slightly lower efficiency compared to CaDDReS-Sc and SeuratCCA (Fig. 5E-F). It is essential to note that CaDDReS-Sc achieved the highest efficiency, predicting responses for all drugs in a single run, while other tools relied on iterative processes. By only focusing on one sample, such as HER2-positive breast cancer sample PM0337, scPharm completed pharmacological subpopulation analysis of single drug within 1 minute across a range of 500 to 2000 cells, displaying significantly lower time consumption compared to all other tools except SeuratCCA (Fig. 5G). Further, we observed runtime stability across multiple repetitions for the sample, whereas CaDDReS-Sc and scDEAL exhibited slight fluctuations in runtime (Fig. 5G). Additionally, the runtime of all tools increased with an escalating number of cells.

### Identification of potential combination regimens

scPharm can be utilized to formulate potential drug combinations using the set covering method, which encompasses booster and compensation effects (Fig. 1). Booster effects involve the use of two distinct drugs with different targets to eliminate a specific subpopulation of tumour cells (Fig. 6A). After applying scPharm to ER-positive breast cancer data, we extracted the top 5 booster effects of ER inhibitions (Fulvestrant and GDC0810) in each sample, along with the combination therapy of other drugs. By investigating these booster effects-based drug combinations, we found that drugs such as Doramapimod and Ribociclib were commonly present in more than 10 samples (out of a total of 14 samples) (Fig. 6B). The correlation of IC50 values of drug pairs in all cancer cell lines was calculated to better understand booster effects between drug pairs. The results showed that both the most frequently occurring Doramapimod and Ribociclib, as well as the less frequent Palbociclib, had a high correlation with ER-targeted drugs, exceeding 0.5 (Fig. 6C). Previous studies have demonstrated the significant tumor regression effects of Fulvestrant and Palbociclib/Ribociclib in ER-positive breast cancer (Fig. 6D) [33]. Ribociclib and Palbociclib are both CDK4/6 inhibitors with similar mechanisms of action and effects, making them clinically applicable for the treatment of ER-positive breast cancer [33, 34]. This strongly supports the credibility of our predicted drug combinations with Fulvestrant or GDC0810. The top-ranked drug, Doramapimod, targets JNK and p38 signaling and is currently only used in clinical trials for psoriasis and rheumatoid arthritis. However, we have reason to speculate that this drug has the potential for adjuvant therapy in ER-positive breast cancer, which would be another possible combination except for the application in acute myeloid leukemia [35].

**Figure 6.**
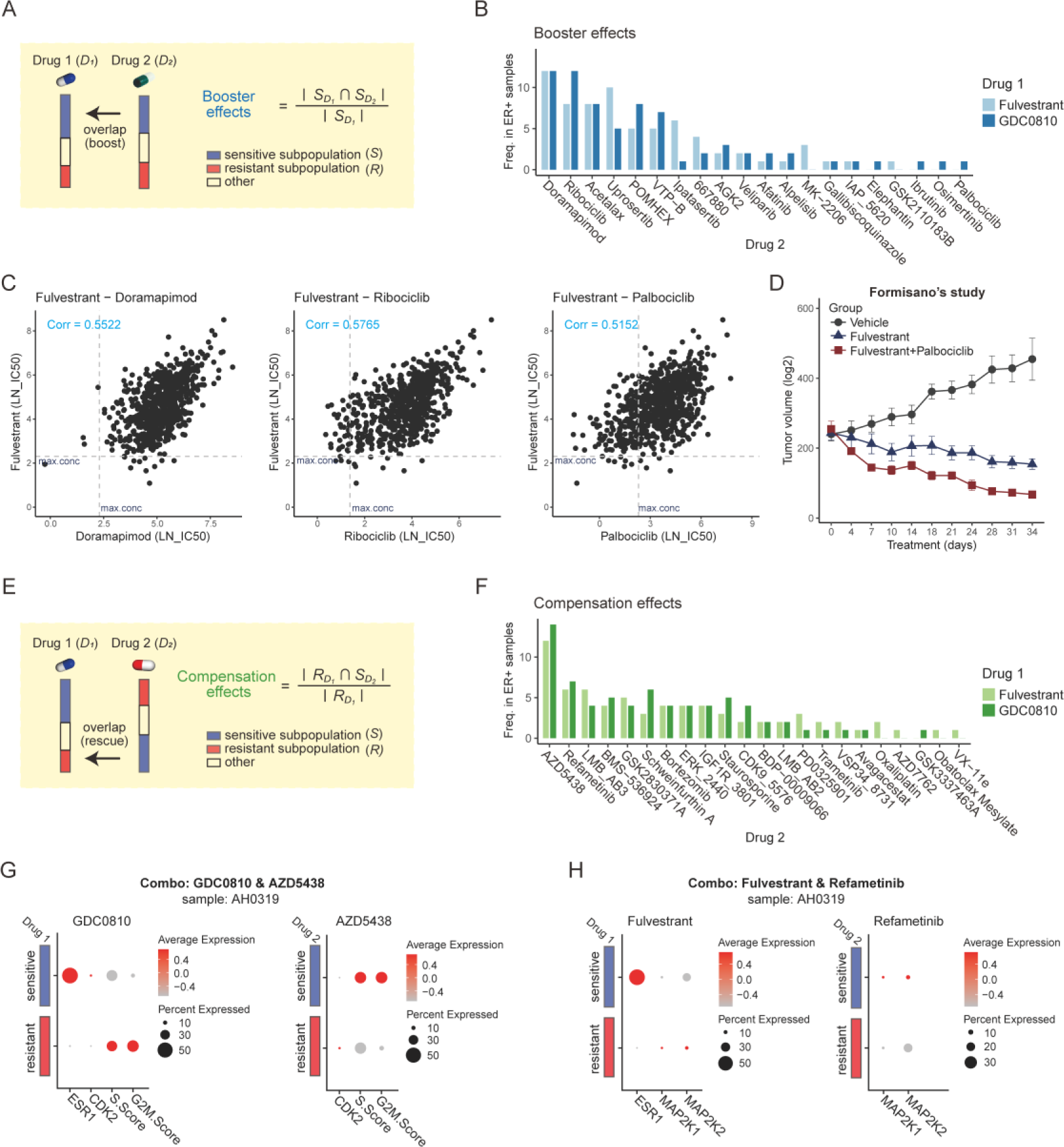
Application of scPharm on drug combinations in ER+ breast cancer. **A.** The booster effects schematic and calculation formula. **B.** Frequency of the top 5 combination medication pairs in each sample predicted by the booster effect across the 14 samples. **C.** Scatterplot of IC50 Pearson correlations for three combination drug pairs in all cancer cell lines. The dashed line indicates the maximum concentration of the drug. **D.** Tumour size diagram of Fulvestrant in combination with Palbociclib at different time points. Data from the study of Formisano et al. **E.** The compensation effects schematic and calculation formula. **F.** Frequency of the top 5 combination medication pairs in each sample predicted by the compensation effects across the 14 samples. **G**. Expression plots of genes and functions indicating drug combination (GDC0810 & AZD5438) in sample AH0319. **H.** Expression plots of genes indicating drug combination (Fulvestrant & Refametinib) in sample AH0319.

Compensation effects suggest that drug 2 can serve as a complementary therapeutic approach to drug 1 (Fig. 6E). Based on the magnitude of compensation effects, the top 5 drug combinations were also extracted for each sample of ER-positive breast cancer. AZD5438, targeting CDK2 [36], appeared in the predicted drug combinations for all samples (Fig. 6F). Analysis revealed significantly higher cell cycle activity in resistant cells to drug 1 (GDC0810) and sensitive cells to drug 2 (AZD5438) (Fig. 6G). This indicates that drug 2 rescues the resistance of drug 1 by inhibiting the ectopic activation of cell cycle in the resistant subpopulation to drug 1. Additionally, Refametinib, a MEK1/2 inhibitor [37], was predicted to have a combined effect in more than 6 samples. Similarly, in Fulvestrant-resistant cells, MAP2K1 and MAP2K2 were significantly upregulated (Fig. 6H). The above results demonstrate the realistic existence of compensation effects and the efficacy of combination pairs predicted by scPharm.

### Prediction of drug side effects on healthy human tissues

scPharm is designed to evaluate drug side effects by utilizing the healthy cells in the tumour microenvironment. In the analysis pipeline of scPharm, the healthy cells were distinguished from tumour cells by CopyKAT method.

To assess side effects, we introduced an index representing the ratio of healthy cells in sensitive subpopulation to all healthy cells (Fig. 7A). Utilizing scPharm, we identified sensitive subpopulation from scRNA-seq data of human healthy tissues, including 5 skin, 8 breast and 12 lung samples, generating a comprehensive landscape of side effects for all drug (Fig. 7B-G). Notably, we observed substantial heterogeneity of side effects both within samples of the same tissue and across different tissues (Fig. 7B, D, F).

**Figure 7.**
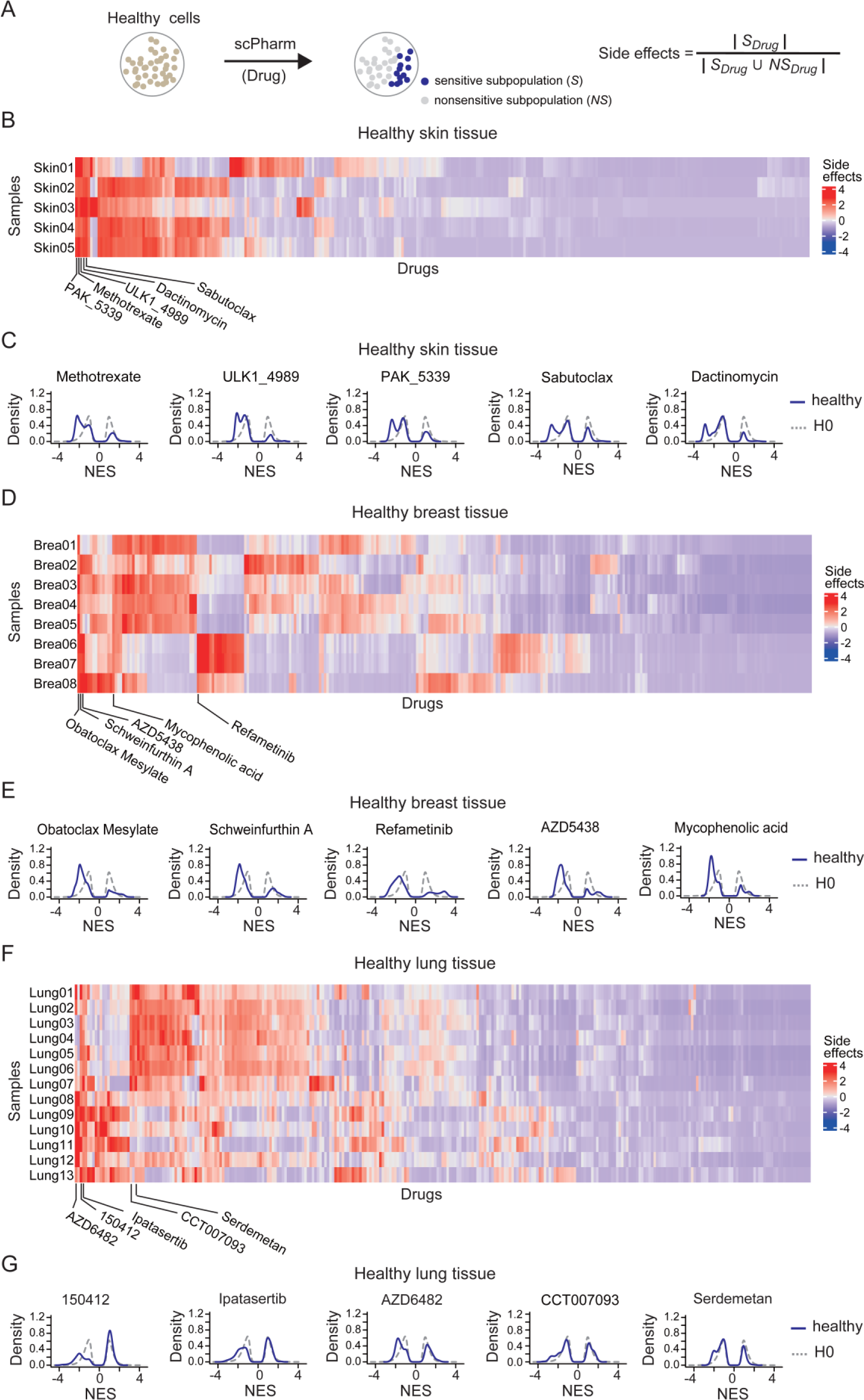
Prediction of drug side effects on healthy human tissues. **A.** Index of side effects evaluation. **B.** Heatmap showing side effects for all drugs in healthy skin tissues from different donors. **C.** Density plot depicting the distributions of NES for the top 5 drugs with the highest index of side effects in pooled healthy skin tissues. **D.** Heatmap showing side effects for all drugs in healthy breast tissues from different donors. **E.** Density plot depicting the distributions of NES for the top 5 drugs with the highest index of side effects in pooled healthy breast tissues. **F.** Heatmap showing side effects for all drugs in healthy lung tissues from different donors. **G.** Density plot depicting the distributions of NES for the top 5 drugs with the highest index of side effects in pooled healthy lung tissues.

For skin tissues, the top 5 drugs with the highest side effects index included Methotrexate, ULK1_4989, PAK_5339, Sabutoclax, and Dactinomycin (Fig. 7B). The NES distribution curves of single healthy cells exhibited dual negative peaks, with one negative peak significantly shifted against H0_sensitive, indicating the presence of a significant sensitive subpopulation to these drugs (Fig. 7C). Methotrexate is an antimetabolite and antifolate drug. It works by inhibiting dihydrofolate reductase, an enzyme involved in the synthesis of DNA, RNA, and proteins. Methotrexate is known to cause various skin-related side effects, including alopecia, radiation recall, reactivation sunburn, and photosensitivity. Dactinomycin, also known as actinomycin D, is a chemotherapy medication used to treat numerous types of cancer. Dactinomycin is known to cause varied skin-related side effects, including rashi, pigment changes, photosensitivity, even more severe skin reactions, such as Stevens-Johnson syndrome and toxic epidermal necrolysis [38, 39].

For breast tissues, the top 5 drugs with the highest side effects index were Obatoclax mesylate, Schweinfurthin A, Refametinib, AZD5438, and Mycophenolic acid (Fig. 7D). The leftward shift of the negative peak against H0_sensitive for these drugs (Fig. 7E) suggests potential toxicity on mammary gland cells. Compared with healthy skin and breast tissues, healthy lung tissues exhibited mild side effects, with even the top 5 drugs showing slightly leftward-shifted negative peaks (Fig. 7F, G), indicating that majority of single cells do not respond significantly to these drugs.

## Discussion

The challenge of molecular heterogeneity within tumours poses a significant obstacle to effective anticancer treatments. Tumours characterized by high intratumoural heterogeneity often lead to unfavorable clinical outcomes, including drug resistance and a poorer prognosis for patients. Thus, the precise identification of tumour heterogeneity becomes crucial for the development of successful therapeutic interventions. Single-cell RNA sequencing (scRNA-seq) technology offers a means to capture gene expression profiles at a single-cell resolution, while drug perturbation experiments yield valuable drug response data at the bulk cell level. Through the integration of these multi-modal datasets, computational methods can effectively transfer drug response information from bulk cells to individual cells. This integration holds the potential to provide a solution for unravelling the complexities of intratumour therapeutic heterogeneity.

In this study, we introduced scPharm, a computational framework designed to integrate large-scale pharmacogenomics profiles with scRNA-seq data. The primary objective was to discern intratumour therapeutic heterogeneity at a single-cell resolution. scPharm assesses the distribution of the identity genes of single cell (Cell-ID) within drug response-determined ranking genes, which is accomplished using the NES statistic obtained from Gene Set Enrichment Analysis (GSEA) as the indicator. Based on NES statistics, scPharm statistically determines sensitive cells and resistant cells in the context of drug perturbation. The framework’s statistical methodology addresses challenges present in current deep-learning models applied in this domain. These challenges include coping with small sample sizes, optimizing label binarization, leveraging data across modalities, and enhancing biological interpretation. Additionally, scPharm goes beyond the prioritization of tailed drugs for a tumour. scPharm stands out with its unique capabilities to predict combination strategies, gauging compensation effects or booster effects between two drugs. Moreover, scPharm is applied to single-cell sequencing data obtained from real tumour tissue scenario. These scenarios involve a mixture of various cell types within the tumour microenvironment, rather than focusing solely on pure cancer cells. By directing attention to these “normal” cells, scPharm holds the potential to evaluate drug toxicity or side effects on non-tumour cells. This capability provides valuable insight into the comprehensive effects of a given drug on the entire tumour tissue.

We conducted a thorough comparative analysis to assess the predictive performance of scPharm against other single-cell prediction tools, including scDEAL and CaDDReS-Sc, as well as label transfer methods such as Scissor and SeuratCCA. While scDEAL employs a deep transfer learning approach to integrate bulk cell line data with scRNA-seq data, while CaDDReS-Sc is a machine learning framework designed for predicting drug responses. It’s worth noting that Scissor and SeuratCCA are not typically used in this domain; Scissor is a method for identifying cell subpopulations associated with a given phenotype, and SeuratCCA is a popular label transfer method within the Seurat package. Our systematic evaluation revealed the superior predictive performance and computational efficiency of scPharm.

Given the advantages of scPharm, there are several points we need to address. First, one key strength of scPharm is rooted in the robust positive correlation between NES statics and AUC values, as depicted in Fig. 2. To reduce the impact of tissue specific expression on prediction, drug resonpse-determined gene lists were generated using the cell lines within the same cancer type. By investigating 12 types of cancer types, we observed the robust correlations consistently present across pan-cancer samples and a diverse array of drugs. The core principle behind applying NES statics to predict drug response relies on calculating NES using single-cell RNA sequencing data and retrieving AUC from the cell population of the same cancer cell lines. While a positive correlation was still observed when NES was calculated based on bulk RNA-seq data, it was not as strong as with single-cell data (Supple. Fig. 7). This may be attributed to the efficiency of the MCA method in extracting Cell-ID from millions of single cells compared to bulk cell samples. Therefore, scPharm takes advantage of data structure of scRNA-seq for improving the correlation between NES statics and drug response.

Second, evaluation of single-cell prediction models at the individual cell level poses challenges since drug response assays are typically conducted on cell populations rather than individual cells. Currently, some publicly sporadic works still use the bulk cells’ data to evaluate the performance of models. To address this, we introduced a novel evaluation approach. Instead of relying on evaluations using bulk cell data, we employed two single-cell RNA-seq datasets, specifically ER-positive and HER2-positive breast cancer tissue samples. These samples, known to be sensitive to ER or HER inhibitions, allowed us to rank all drugs based on their corresponding drug sensitivity. The ranking of ER or HER inhibitions at the top positions indicates good prediction performance. This innovative approach is a step forward in evaluating the prediction performance of single-cell samples, especially when drug response labels are not readily available.

Finally, while the current version of scPharm excels in predict drug toxicity or side-effects on “normal” cells in tumour tissue, its potential can be extended to a large-scale investigation based on large samples as well as the precise classifications of “normal” cells in the tumour microenvironment, such as T cell, B cells and fibroblasts. This expansion will enable a more comprehensive understanding of the impact of drugs on both the tumour itself and its microenvironment.

In summary, scPharm is a computational framework tailored for scRNA-seq data, integrating pharmacogenomics profiles to uncover therapeutic heterogeneity within tumours at single-cell resolution. The tool not only prioritizes tailored drugs but also provides insight into combination therapy regimen and drug toxicity in cancers.

## Materials and methods

### Data source

The pharmacogenomic dataset utilized in this study encompasses 969 human cancer cell lines, 286 drugs, and a total of 242,036 drug response measurements. The bulk RNA-seq data were sourced from Cell Model Passports database (https://cellmodelpassports.sanger.ac.uk/). Drug response data were obtained from the GDSC2 project of the GDSC database (https://www.cancerrxgene.org/. release 8.4, July 2022) [27]. The study quantifies drug response in terms of Area Under the dose response Curve (AUC) and half maximal inhibitory concentration (IC50), where higher values indicate drug resistance and lower values denote drug sensitivity in cancer cell lines. The scRNA-seq data of human cancer cell lines were collected from the SCP542 dataset on Single Cell Portal (https://singlecell.broadinstitute.org/single_cell). Single-cell data from a total of 14 cancers were used in this study.

To evaluate our study, the scRNA-seq data were obtained from the NCBI GEO database, specifically from GSE161529[13], GSE158677[15] and GSE134839[14]. These datasets respectively include scRNA-seq profiles for 6 HER-positive and 14 ER-positive breast cancer tumours, 5 mice from MMTV-PyMT mouse mammary tumour model, and lung adenocarcinoma cell line PC9 treated with Erlotinib at Day 1, Day 2, Day 4, Day 9 and Day 11.

To model a statistical test, scRNA-seq data from GSE196638, GSE122960, GSE150247, GSE164898 and GSE151177 were used to construct null distributions (H0) of the test. These datasets include scRNA-seq profile from healthy human tissues of lung [21-23], skin [24], and breast [25]. The datasets were also utilized to evaluate drug side effects. To calculate pathway activity, we collected canonical pathways from the KEGG database. All source data are provided with this article.

### Data processing

For the bulk RNA-seq data, gene expression profiles were represented using the normalized expression index - Transcripts Per Million (TPM), with the removal of missing values and genes that showed no expression across all cell lines. Notably, only cell lines with corresponding drug response information were retained for analysis. In the case of scRNA-seq data, the R package Seurat (v4.3.0) [40] was employed for data processing, including reading and initial quality control. Quality control was conducted by adjusting parameters such as maxGene and maxUMI based on the expression profile characteristics of the samples, followed by normalization and dimensionality reduction clustering.

### The framework of scPharm

scPharm is a comprehensive framework composed of two interconnected modules designed for the analysis and prediction of pharmacological subtypes of single cells, as well as prioritization of the tailored drugs and exploration of combinatorial drug usage and potential drug side effects (Fig. 1).

<u>Module 1 of scPharm</u> utilizes gene expression profiles from cancer cell lines measured by bulk RNA-seq and their corresponding drug response data from Cell Model Passports database and GDSC database. It applies the Pearson correlation method to calculate the correlations between gene expression and drug response, and generates drug response-determined gene list for each drug within a specific cancer type. Genes at the top of the ranked list, whose expression positively correlated with AUC, are linked to drug resistance, while those at the bottom are associated with drug sensitivity.

In the real scenario where scRNA-seq data is derived from tumour tissues rather than cancer cell lines, scPharm employs CopyKAT to differentiate between cancer cells and healthy cells based on gene copy number variations[19]. Cancer cells are used for identifying pharmacological subpopulations and rational medicine, while healthy cells are employed to assess potential drug side effects. Subsequently, the Multiple Correspondence Analysis (MCA) method is utilized to extract gene signature from individual cells, effectively defining as single-cell identity (Cell-ID) [20]. Leveraging drug response-determined gene list and Cell-ID as inputs, the method Gene Set Enrichment Analysis (GSEA) [12] evaluates whether a single cell is sensitive to a drug or not, depending on the distribution of its Cell-ID throughout the ranked gene list for the specific drug. If the Cell-ID of an individual cell is predominantly enriched at the top of the list, the cell is predicted to be resistant to the tested drug, while enrichment at the bottom suggests sensitivity. The normalized enrichment score (NES) obtained through GSEA serves as a statistic indicating the drug response of a single cell, categorizing it as sensitive, resistant, or other.

To model the null distribution of NES in the context of drug response, scRNA-seq data from healthy human tissues of lung, skin and breast were collected [21-25]. Each tissue sample was integrated into a single-cell object using Seurat. Subsequently, we conducted quality control, normalization, dimensionality reduction, clustering, MCA and GSEA analysis on each tissue sample. After completion, the NES statistics for all cells related to each drug were merged to serves as the null distribution for this study. Gaussian mixture model decomposition is performed using the normalmixEM method in the Mixtools package to accurately estimate the parameter of the null distribution [26]. Using a unified threshold based on the mean plus or minus one standard deviation, scPharm categorizes tumour single-cell data into sensitive subpopulation or resistant subpopulation for a given drug, enhancing the interpretability and reliability of the framework.

<u>Module 2 of scPharm</u> evaluates the suitability of drugs within a specific tumour sample by defining a score, denoted as ‘Dr’, based on the number of sensitive and resistant single cells obtained from Module 1. The score is computed as *Dr* = *R*_*s*_ ∗ (1 − *R*_*r*_), where *R*_*s*_ represents the ratio of sensitive subpopulation and *R*_*r*_ represents the ratio of resistant subpopulation in the tumour sample. Based on this score, scPharm ranks drugs to provide drug recommendations.

scPharm also explores potential combinatorial drug therapies using compensation and booster effects. Compensation involves the second drug sensitizing drug-resistant cells that are unresponsive to the first drug, thereby compensating for the lack of response. In contrast, booster effects occur when the second drug synergistically enhances the cytotoxic effect of the first drug on sensitive tumour cells.

Set covering method is employed to quantify the combination effects of two drugs described above.

For compensation effects, the effect is calculated with this formula,

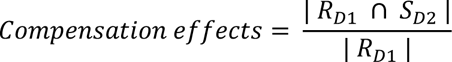

For booster effects, the effect is calculated with this formula,

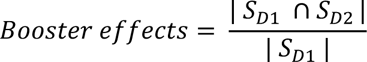

where *R*_*D*1_ represents the resistant cell subpopulation to Drug 1, *S*_*D*2_ represents the sensitive cell subpopulation to Drug 2, and *S*_*D*1_ represents the sensitive cell subpopulation to Drug 1.

Additionally, scPharm considers healthy cells infiltrating tumour microenvironment in its analysis. If a drug is found to be also sensitive to these cells, it indicates its potential toxicity to healthy cells and is classified as having side effects. To quantify the side effects of a drug on a specific patient, we constructed the index described by this formula,

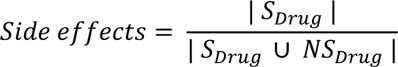

where the *S*_*Drug*_ represents the sensitive cell subpopulation to *Drug* in healthy cells and *NS*_*Drug*_ represents the nonsensitive cell subpopulation to *Drug*.

### Evaluation of scPharm

In the evaluation of scPharm, two tumour single-cell drug sensitivity prediction tools were employed: scDEAL[10] and CaDDReS-Sc[16]. Label transfer methods, Scissor [17] and SeuratCCA[18], were also utilized to transfer the response label of pharmacogenomic data to tumour single-cell data.

scDEAL is a deep transfer learning approach that integrates bulk cell line data with scRNA-seq data to predict cancer drug responses at the single-cell level. In our evaluation, scDEAL code was executed with default parameters on scRNA-seq data and drugs, and the predicted drug response in scDEAL, sens_label (1: sensitive, 0: resistant) was extracted for evaluation.

CaDDReS-Sc is a machine learning framework that combines scRNA-seq data with matrix factorization techniques to predict drug responses in heterogeneous tumours. In our evaluation, drug sensitivity in single cells was predicted using default parameters, and the calculated cell death rate (cell_death) for each drug was extracted as the ranking result for evaluation. Considering that CaDDReS-Sc defaults to using GDSC1 data, we additionally expanded the training and prediction of CaDDReS-Sc using GDSC2 data to facilitate result comparison (named CaDDReS-Sc.gdsc2).

Scissor is a method used to identify cell subpopulations associated with a given phenotype by leveraging single-cell data. It quantifies the similarity between individual cells and bulk samples, integrating phenotype-associated bulk expression data with single-cell data. In our evaluation, Scissor was employed to transfer labels from pharmacogenomic data to scRNA-seq data. For each drug-treated bulk cancer cell line data, the cell lines were first divided into sensitive and resistant cells based on the maximum drug concentration using IC50 values, which served as the phenotypic input for Scissor. Subsequently, Scissor was utilized in binomial mode to accomplish the prediction of tumour cells.

SeuratCCA, a label transfer method in the Seurat package, allows integration of single-cell measurements across different scRNA-seq technologies and modalities. Bulk cancer cell line data was binarized into sensitive and resistant labels, based on the maximum drug concentration using IC50 values. The FindTransferAnchors method was then used to compute anchors between the cancer cell line expression profiles and scRNA-seq data. Finally, the MapQuery method was used to transfer the bulk cancer cell-line labels to the scRNA-seq data.

Given the lack of gold standard data for predicting single-cell drug responses using traditional benchmark methods, we introduced a novel evaluation approach focused on comparing the drug recommendations generated by different tools. With the exception of CaDDReS-Sc, which calculates drug recommendation rankings based on the cancer cell kill scores, the remaining tools calculate drug recommendation rankings using Dr scores defined in scPharm. The systematic comparisons were evaluated on scRNA-seq data of HER2-positive and ER-positive breast cancer samples[13]. These samples are known to be sensitive to HER inhibitions or ER inhibitions. All evaluations were conducted using the same system configuration, with a CPU core limit of 20.

### Statistical analysis

To compare NES statistics between sensitive group and resistant group classified by AUC values, we conducted a one-sided Mann-Whitney test [41]. If the NES statistics in resistant group are greater than sensitive group and p-value is less than 0.1, we conclude there is a significant difference.

To compare pathway activity, the significance of differences in pathway activity between HER2-positive breast cancer samples is assessed using Mann-Whitney test. The symbols “ns”, “*”, “**”, “***” and “****” represent the significance levels as follows: 0.05 ≤ p < 1, 0.01 ≤ p < 0.05, 0.001 ≤ p < 0.01, p < 0.001, p < 0.0001, respectively.

To compare single-cell subpopulations of the same drug with the different maximum screening concentrations, we conducted a one-sided Mann-Whitney test. The symbols “ns”, “*”, “**” and “***” represent the significance levels as follows: 0.05 ≤ p < 1, 0.01 ≤ p < 0.05, 0.001 ≤ p < 0.01, p < 0.001, respectively.

## Authors’ contributions

Haiyun Wang conceived the hypothesis. Peng Tian, Jie Zheng and Haiyun Wang designed and performed the data analysis. Yue Xu, Tao Wu, Shuting Chen, Yinuo Zhang, Bingyue Zhang, Keying Qiao, and Yuxiao Fan collected and preprocessed the data. Chiara Ambrogio participated in the helpful discussions. Peng Tian, Jie Zheng and Haiyun Wang interpreted the results and wrote the manuscript.

## Data Availability

The datasets generated and/or analysed during the current study are available in the Gene Expression Omnibus (GEO) [https://www.ncbi.nlm.nih.gov/geo/] and Single Cell Portal (SCP) [https://singlecell.broadinstitute.org/single_cell]

## Code Availability

scPharm is implemented in R and is freely available from GitHub (https://github.com/WangHYLab/scPharm).

---

**Supple Figure 1.**
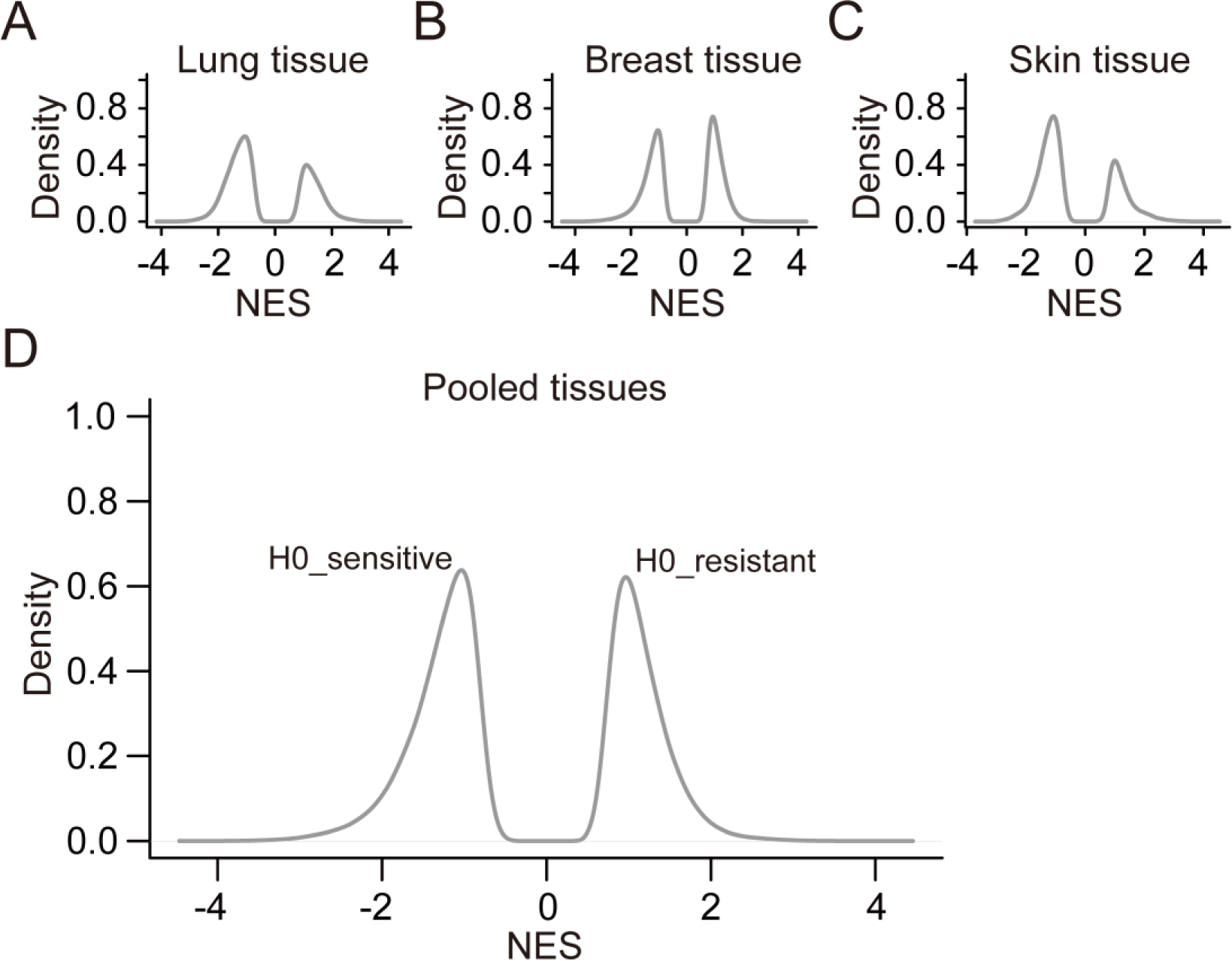
Null distribution of NES. **A.** Density plot depicting the NES statistics of single cells from healthy lung tissues across all drugs. **B.** Density plot depicting the NES statistics of single cell from healthy breast tissues across all drugs. **C.** Density plot depicting the NES statistics of single cells from healthy skin tissues across all drugs. **D.** Density plot depicting the NES statistics of single cells from pooled healthy tissues across all drugs.

**Supple Figure 2.**
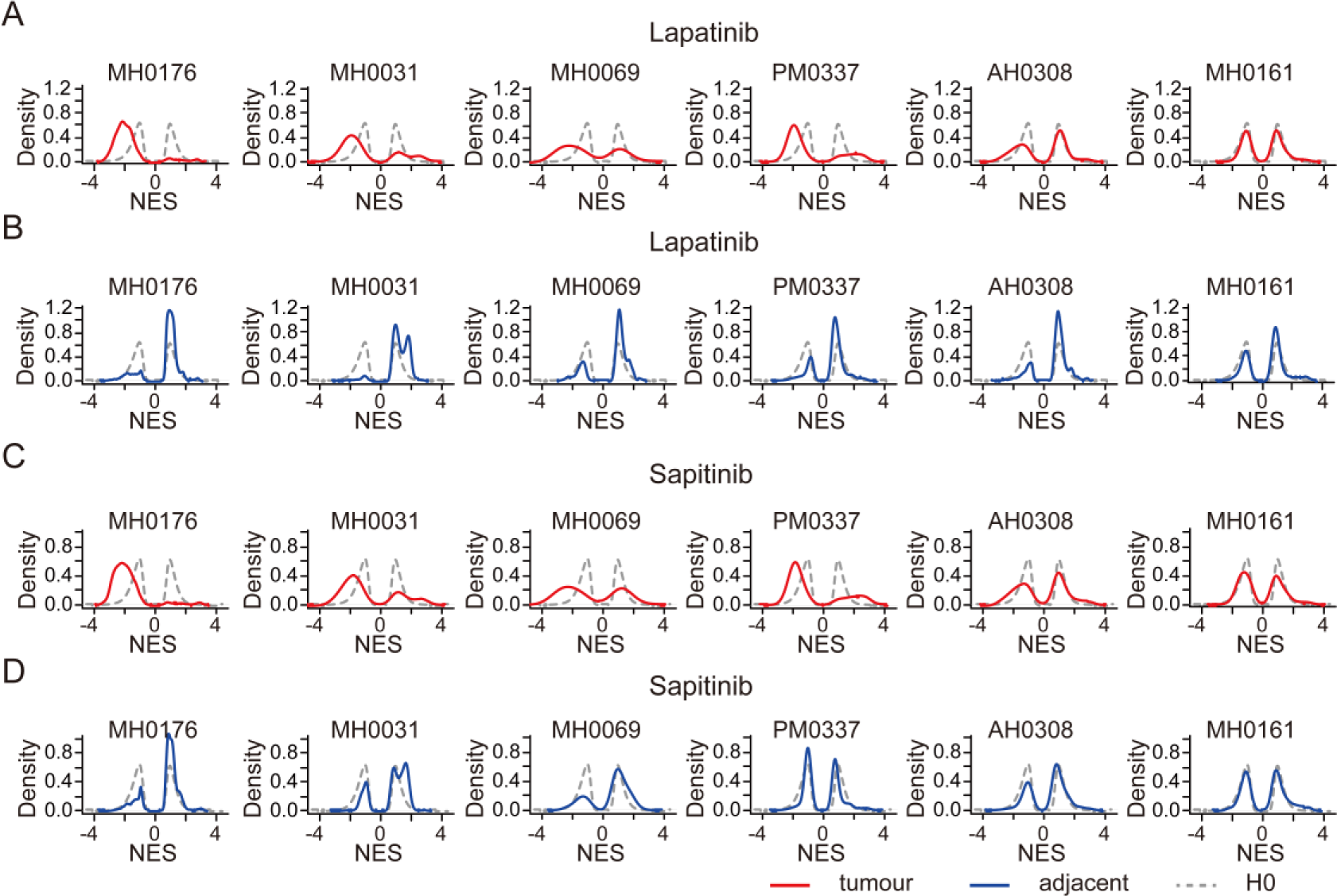
scPharm is applied in HER2+ breast cancer. **A.** Density plot depicting the NES statistics of single cells from six HER2+ breast cancer tissues (red curves) and from healthy human tissues (grey curves) in the context of Lapatinib. **B.** Density plot depicting the NES statistics of single cells from tumour-adjacent tissues and from healthy human tissues in the context of Lapatinib. **C.** Density plot depicting the NES statistics of single cells from six HER2+ breast cancer tissues (red curves) and from healthy human tissues (grey curves) in the context of Sapitinib. **D.** Density plot depicting the NES statistics of single cells from tumour-adjacent tissues and from healthy human tissues in the context of Sapitinib.

**Supple Figure 3.**
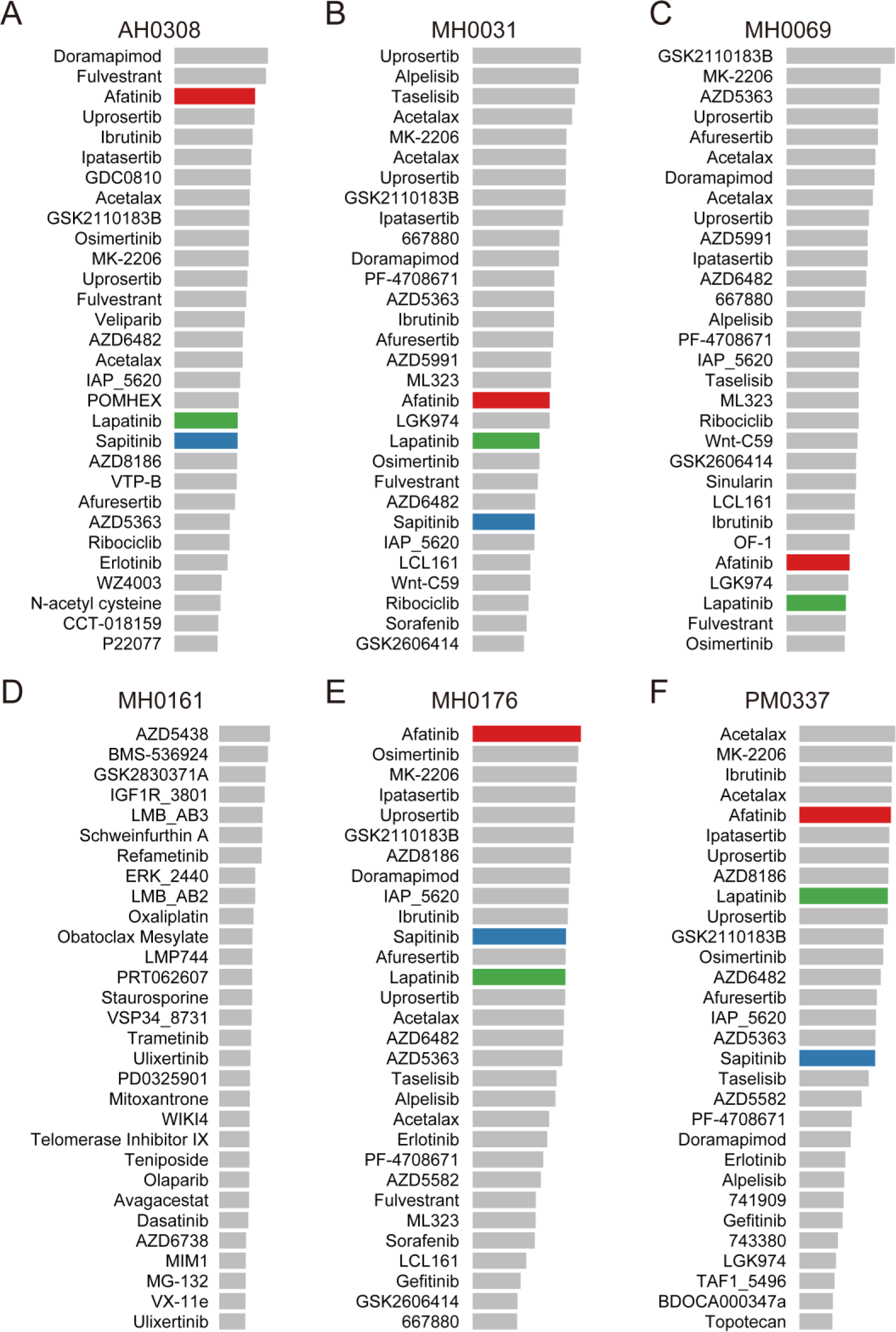
The recommended drugs for HER2+ samples. **A-F.** The top 30 recommended drugs for AH0308, MH0031, MH0069, MH0161, MH0176 and PM0337 respectively. The length of the bars represents the level of effectiveness.

**Supple Figure 4.**
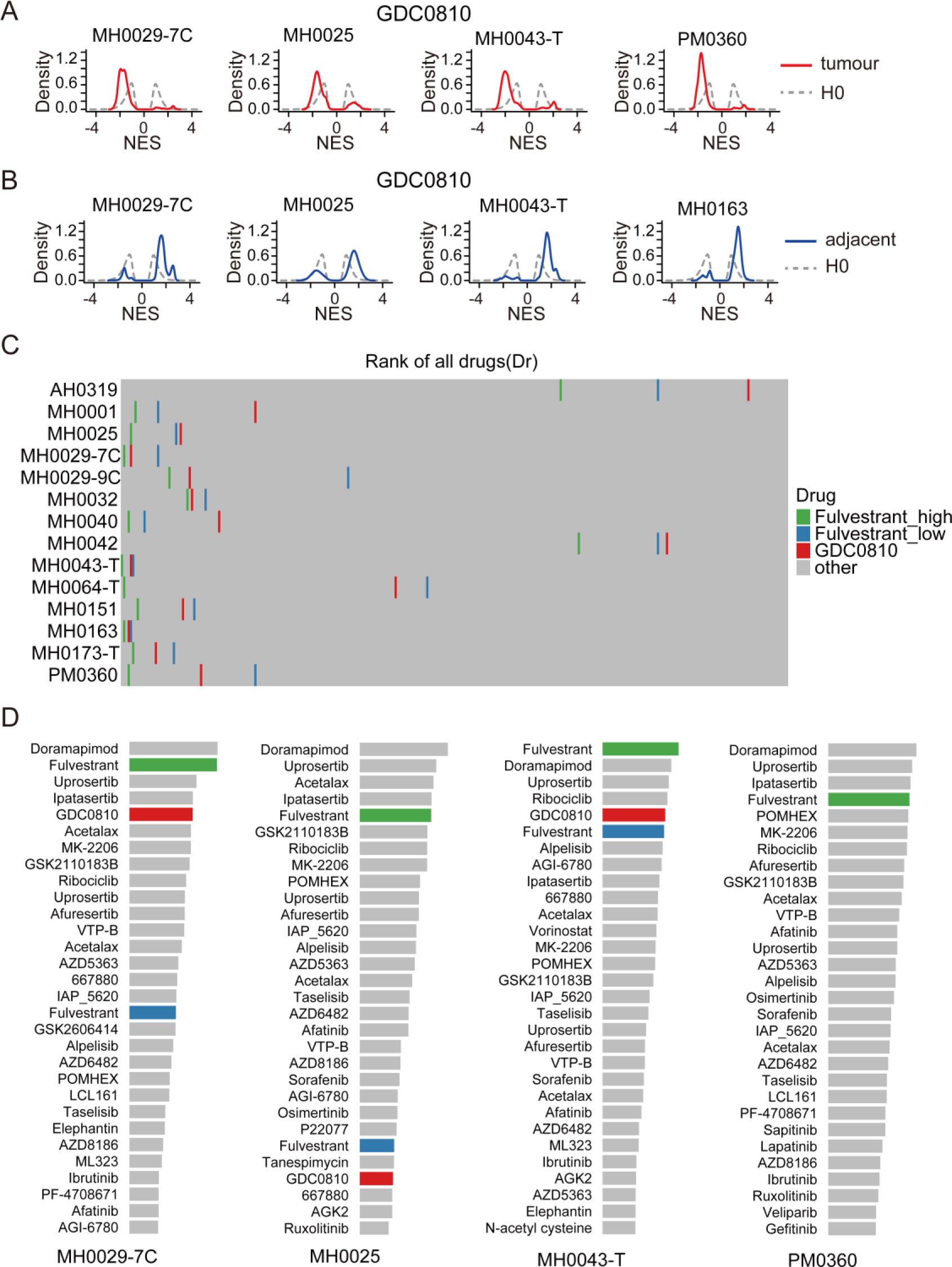
scPharm is applied in ER+ breast cancer. **A.** Density plot depicting the NES statistics of single cells from four ER+ breast cancer tissues (red curves) and from healthy human tissues (grey curves) in the context of GDC0810. **B.** Density plot depicting the NES statistics of single cells from tumour-adjacent tissues and from healthy human tissues in the context of GDC0810. **C.** Rank of two ER inhibitions in ER+ samples. **D.** The top 30 recommended drugs for MH0029-7C, MH0025, MH0069, MH0043-T and PM0360 respectively. The length of the bars represents the level of effectiveness.

**Supple Figure 5.**
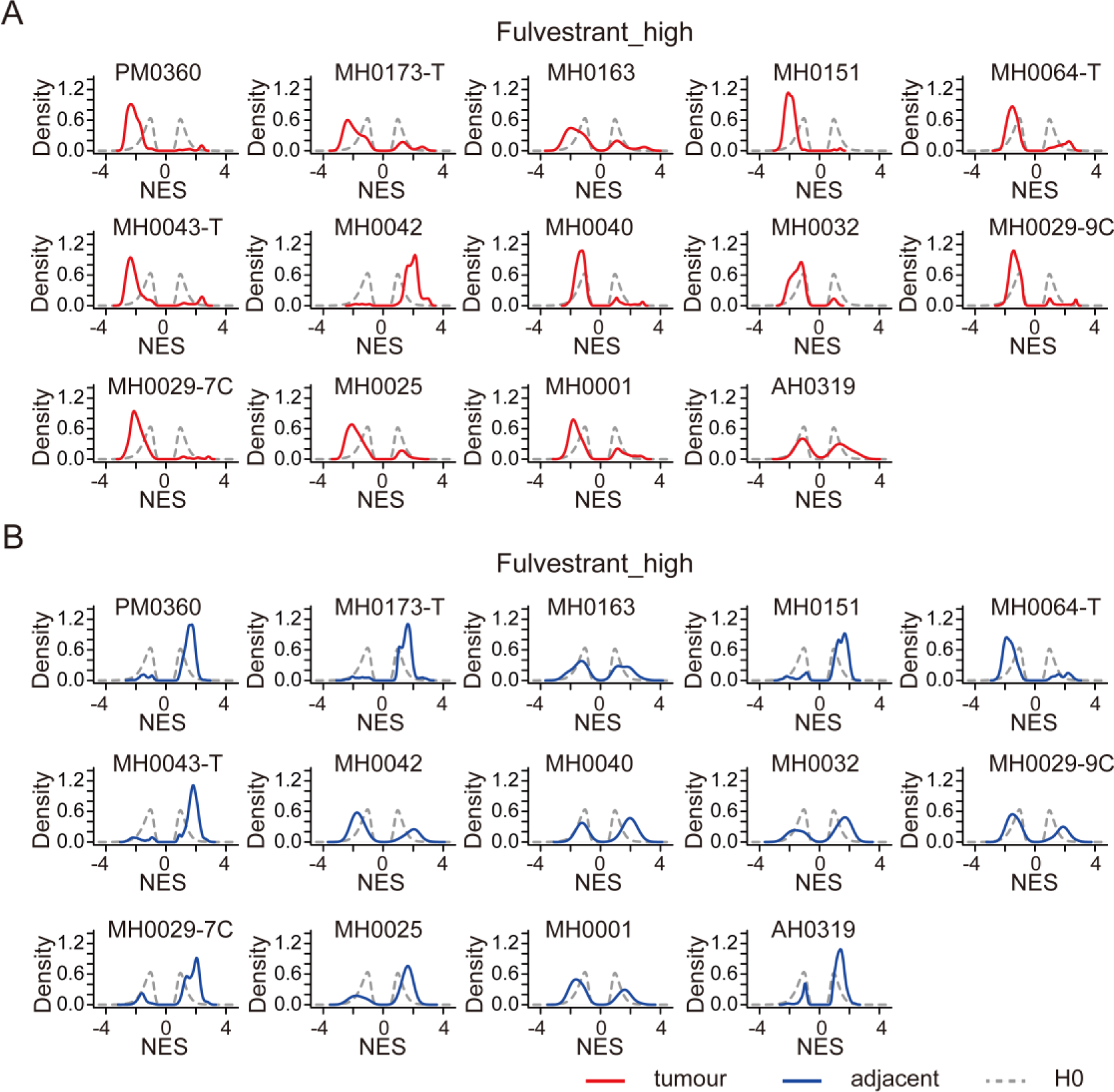
scPharm is applied in ER+ breast cancer in the context of Fulvestrant. **A.** Density plot depicting the NES statistics of single cells from fourteen ER+ breast cancer tissues (red curves) and from healthy human tissues (grey curves) in the context of Fulvestrant_high. **B.** Density plot depicting the NES statistics of single cells from tumour-adjacent tissues and from healthy human tissues in the context of Fulvestrant_high.

**Supple Figure 6.**
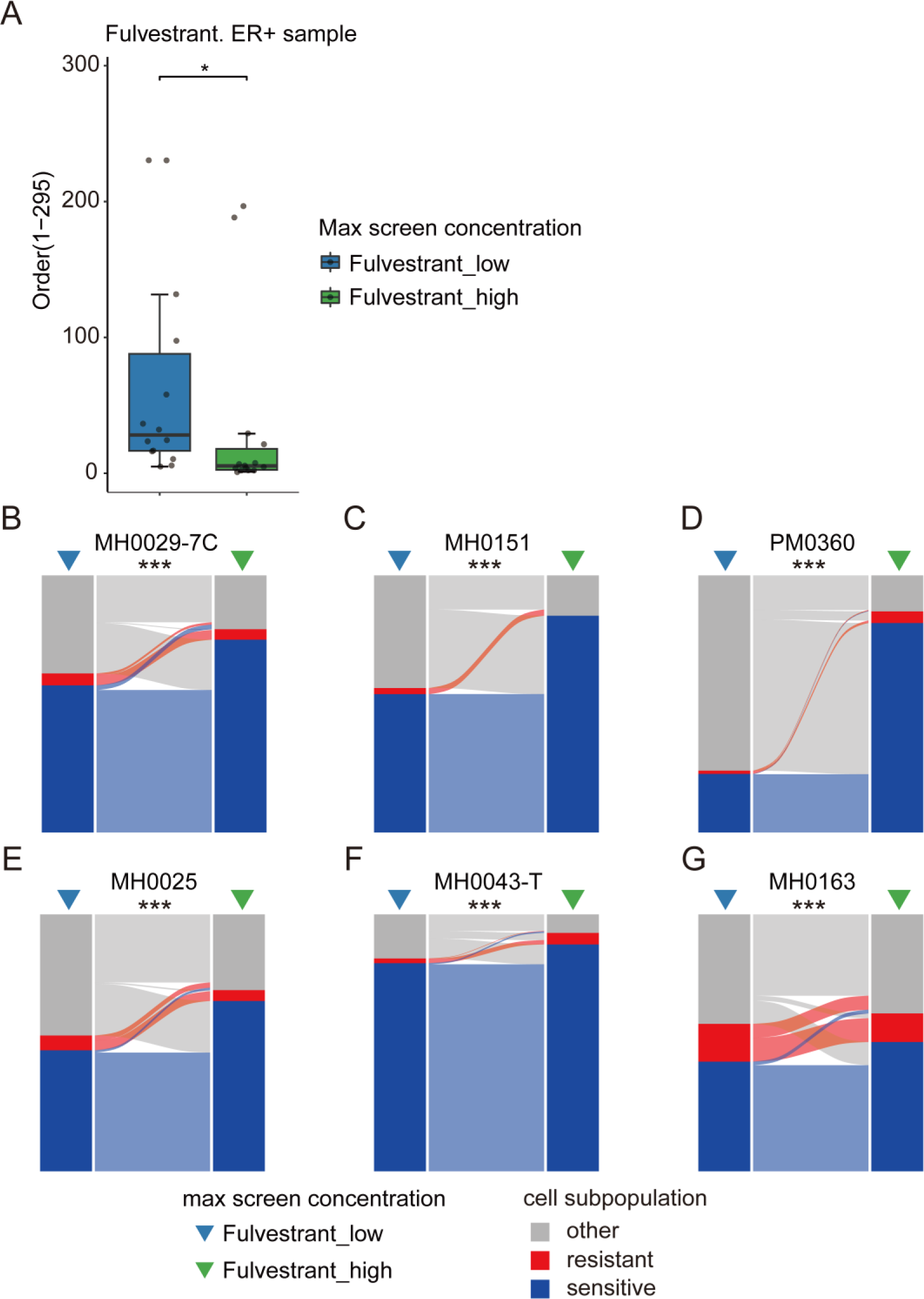
Differences at the different maximum testing concentrations. **A.** Comparative ranking of predictions for Fulvestrant at various concentrations. **B-G.** Alluvial plot showing the changes of three types of subpopulations between the different concentrations across the samples. The Wilcoxon rank-sum test was employed, with “*” indicating a p-value less than 0.05, “**” indicating a p-value less than 0.01, and “***” indicating a p-value less than 0.001.

**Supple Figure 7.**
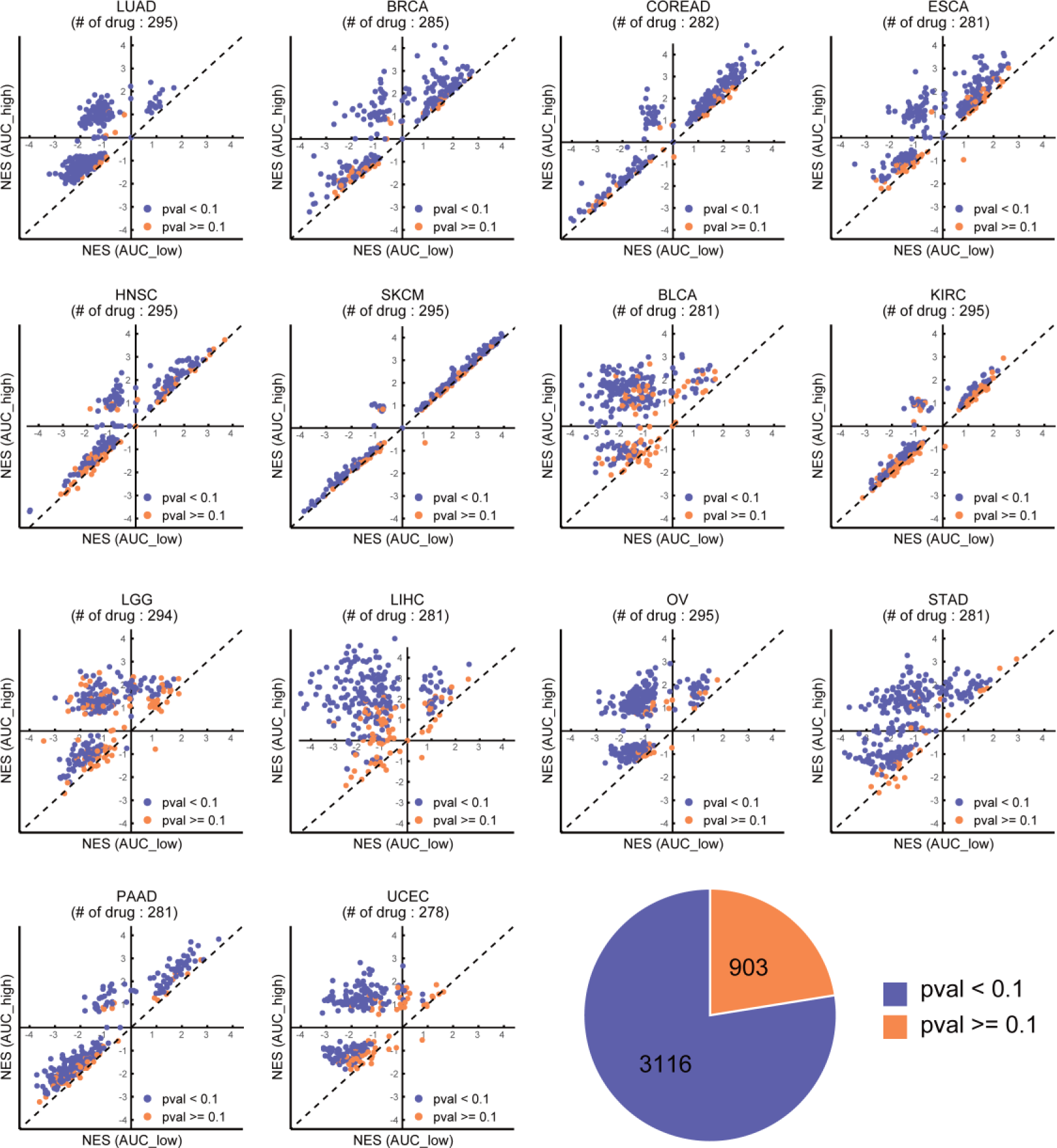
The difference of cell line’s NES between various groups. Scatter plot showing the NES statistics in the AUC-high group versus the AUC-low group. Cell lines are divided into AUC-high and AUC-low groups based on the median of AUC values to each drug.

## References

[1] Dixit A, Parnas O, Li B et al. Perturb-Seq: Dissecting Molecular Circuits with Scalable Single-Cell RNA Profiling of Pooled Genetic Screens. Cell 2016; 167:1853–1866 e1817.

[2] Papalexi E, Mimitou EP, Butler AW et al. Characterizing the molecular regulation of inhibitory immune checkpoints with multimodal single-cell screens. Nat Genet 2021; 53:322–331.

[3] Srivatsan SR, McFaline-Figueroa JL, Ramani V et al. Massively multiplex chemical transcriptomics at single-cell resolution. Science 2020; 367:45–51.

[4] Barretina J, Caponigro G, Stransky N et al. The Cancer Cell Line Encyclopedia enables predictive modelling of anticancer drug sensitivity. Nature 2012; 483:603–607.

[5] Garnett MJ, Edelman EJ, Heidorn SJ et al. Systematic identification of genomic markers of drug sensitivity in cancer cells. Nature 2012; 483:570–575.

[6] Rees MG, Seashore-Ludlow B, Cheah JH et al. Correlating chemical sensitivity and basal gene expression reveals mechanism of action. Nat Chem Biol 2016; 12:109–116.

[7] Behan FM, Iorio F, Picco G et al. Prioritization of cancer therapeutic targets using CRISPR-Cas9 screens. Nature 2019; 568:511–516.

[8] Meyers RM, Bryan JG, McFarland JM et al. Computational correction of copy number effect improves specificity of CRISPR-Cas9 essentiality screens in cancer cells. Nat Genet 2017; 49:1779–1784.

[9] McDonald ER, 3rd, de Weck A, Schlabach MR et al. Project DRIVE: A Compendium of Cancer Dependencies and Synthetic Lethal Relationships Uncovered by Large-Scale, Deep RNAi Screening. Cell 2017; 170:577–592 e510.

[10] Chen JY, Wu ZY, Qi R et al. Deep transfer learning of cancer drug responses by integrating bulk and single-cell RNA-seq data. Nature Communications 2022; 13.

[11] Suphavilai C, Chia S, Sharma A et al. Predicting heterogeneity in clone-specific therapeutic vulnerabilities using single-cell transcriptomic signatures. Genome Med 2021; 13:189.

[12] Subramanian A, Tamayo P, Mootha VK et al. Gene set enrichment analysis: A knowledge-based approach for interpreting genome-wide expression profiles. P Natl Acad Sci USA 2005; 102:15545–15550.

[13] Pal B, Chen YS, Vaillant F et al. A single-cell RNA expression atlas of normal, preneoplastic and tumorigenic states in the human breast. Embo Journal 2021; 40.

[14] Aissa AF, Islam A, Ariss MM et al. Single-cell transcriptional changes associated with drug tolerance and response to combination therapies in cancer. Nat Commun 2021; 12:1628.

[15] Valdes-Mora F, Salomon R, Gloss BS et al. Single-cell transcriptomics reveals involution mimicry during the specification of the basal breast cancer subtype. Cell Rep 2021; 35:108945.

[16] Suphavilai C, Chia SM, Sharma A et al. Predicting heterogeneity in clone-specific therapeutic vulnerabilities using single-cell transcriptomic signatures. Genome Medicine 2021; 13.

[17] Sun DC, Guan XN, Moran AE et al. Identifying phenotype-associated subpopulations by integrating bulk and single-cell sequencing data. Nature Biotechnology 2022; 40:527-+.

[18] Butler A, Hoffman P, Smibert P et al. Integrating single-cell transcriptomic data across different conditions, technologies, and species. Nat Biotechnol 2018; 36:411–420.

[19] Gao RL, Bai SS, Henderson YC et al. Delineating copy number and clonal substructure in human tumors from single-cell transcriptomes. Nat Biotechnol 2021; 39:599–608.

[20] Cortal A, Martignetti L, Six E, Rausell A. Gene signature extraction and cell identity recognition at the single-cell level with Cell-ID. Nat Biotechnol 2021; 39:1095-+.

[21] Wang CQ, Hyams B, Allen NC et al. Dysregulated lung stroma drives emphysema exacerbation by potentiating resident lymphocytes to suppress an epithelial stem cell reservoir. Immunity 2023; 56:576-+.

[22] Reyfman PA, Walter JM, Joshi N et al. Single-Cell Transcriptomic Analysis of Human Lung Provides Insights into the Pathobiology of Pulmonary Fibrosis. American Journal of Respiratory and Critical Care Medicine 2019; 199:1517–1536.

[23] Kathiriya JJ, Wang CQ, Zhou MQ et al. Human alveolar type 2 epithelium transdifferentiates into metaplastic KRT5 basal cells. Nature Cell Biology 2022; 24:10-+.

[24] Kim J, Lee J, Kim HJ et al. Single-cell transcriptomics applied to emigrating cells from psoriasis elucidate pathogenic versus regulatory immune cell subsets. J Allergy Clin Immunol 2021; 148:1281–1292.

[25] Bhat-Nakshatri P, Gao H, Sheng L et al. A single-cell atlas of the healthy breast tissues reveals clinically relevant clusters of breast epithelial cells. Cell Rep Med 2021; 2:100219.

[26] Benaglia T, Chauveau D, Hunter DR, Young DS. mixtools: An R Package for Analyzing Finite Mixture Models. J Stat Softw 2009; 32:1–29.

[27] Yang WJ, Soares J, Greninger P et al. Genomics of Drug Sensitivity in Cancer (GDSC): a resource for therapeutic biomarker discovery in cancer cells. Nucleic Acids Research 2013; 41:D955–D961.

[28] Britten CD, Finn RS, Bosserman LD et al. A phase I/II trial of trastuzumab plus erlotinib in metastatic HER2-positive breast cancer: a dual ErbB targeted approach. Clin Breast Cancer 2009; 9:16–22.

[29] Yoldi G, Pellegrini P, Trinidad EM et al. RANK Signaling Blockade Reduces Breast Cancer Recurrence by Inducing Tumor Cell Differentiation. Cancer Res 2016; 76:5857–5869.

[30] Mohr CJ, Gross D, Sezgin EC et al. KCa3.1 Channels Confer Radioresistance to Breast Cancer Cells. Cancers (Basel) 2019; 11.

[31] Varticovski L, Hollingshead MG, Robles AI et al. Accelerated preclinical testing using transplanted tumors from genetically engineered mouse breast cancer models. Clin Cancer Res 2007; 13:2168–2177.

[32] Chen Q, Liu X, Luo Z et al. Chloride channel-3 mediates multidrug resistance of cancer by upregulating P-glycoprotein expression. J Cell Physiol 2019; 234:6611–6623.

[33] Formisano L, Lu Y, Servetto A et al. Aberrant FGFR signaling mediates resistance to CDK4/6 inhibitors in ER+ breast cancer. Nat Commun 2019; 10:1373.

[34] Braal CL, Jongbloed EM, Wilting SM et al. Inhibiting CDK4/6 in Breast Cancer with Palbociclib, Ribociclib, and Abemaciclib: Similarities and Differences. Drugs 2021; 81:317–331.

[35] Kurtz SE, Eide CA, Kaempf A et al. Associating drug sensitivity with differentiation status identifies effective combinations for acute myeloid leukemia. Blood Adv 2022; 6:3062–3067.

[36] Boss DS, Schwartz GK, Middleton MR et al. Safety, tolerability, pharmacokinetics and pharmacodynamics of the oral cyclin-dependent kinase inhibitor AZD5438 when administered at intermittent and continuous dosing schedules in patients with advanced solid tumours. Ann Oncol 2010; 21:884–894.

[37] Subbiah V, Baik C, Kirkwood JM. Clinical Development of BRAF plus MEK Inhibitor Combinations. Trends Cancer 2020; 6:797–810.

[38] Wang S, Li T, Wang Y et al. 5-Fluorouracil and actinomycin D lead to erythema multiforme drug eruption in chemotherapy of invasive mole: Case report and literature review. Medicine (Baltimore) 2022; 101:e31678.

[39] Liang Y, Yang Z, Xu ZG, Ma L. Toxic epidermal necrolysis after dactinomycin and vincristine combination chemotherapy for nephroblastoma. J Zhejiang Univ Sci B 2017; 18:649–652.

[40] Hao YH, Hao S, Andersen-Nissen E et al. Integrated analysis of multimodal single-cell data. Cell 2021; 184:3573-+.

[41] Bauer DF. Constructing Confidence Sets Using Rank Statistics. Journal of the American Statistical Association 1972; 67:687–690.

